# The *Orshina* rhythm in a colonial urochordate: recurrent aging/rejuvenation sequels

**DOI:** 10.1101/2023.01.30.526091

**Authors:** Oshrat Ben-Hamo, Ido Izhaki, Rachel Ben-Shlomo, Baruch Rinkevich

**Affiliations:** National Institute of Oceanography, Tel Shikmona, P.O. Box 9753, Haifa 3109701, Israel; Department of Evolutionary and Environmental Biology, Faculty of Natural Sciences, University of Haifa, Mount Carmel, Haifa 3498838, Israel; Department of Biology and Environment, Faculty of Natural Sciences, University of Haifa – Oranim, Tivon 36006, Israel

## Abstract

When it comes to aging, some colonial invertebrates present disparate patterns from the customary aging phenomenon in unitary organisms, where a single senescence phenomenon along ontogeny culminates in their inevitable deaths. Here we studied aging processes in 81 colonies of the marine urochordate *Botryllus schlosseri* each followed from birth to death (over 720 days). The colonies were divided between three life history strategies, each distinct from the others based on the existence/absence of colonial fission: NF (no fission), FA (fission develops after the colony reaches maximal size), and FB (fission develops before the colony reaches maximal size). Results revealed that sexual reproductive statuses (hermaphroditism and male only settings), colonial vigorousness and sizes, represent coinciding and repeated rhythms of one or more emerged life/death ‘astogenic segments’ on the whole-genet level, each is termed as *Orshina*, and the sum of all segments as the *Orshina* rhythm. Each *Orshina* segment lasts about three months (containing ca. 13 blastogenic cycles), ends by either the colonial death or rejuvenation, and manipulated by absence/existing of fission events in NF/FA/FB strategies. These findings indicate that reproduction, life span, death, rejuvenation and fission events are important scheduled biological components in the constructed *Orshina* rhythm, a novel aging phenomenon.

## Introduction

The nature of life does not prohibit an indefinite, long lasting life span of multicellular organisms. Some organisms may escape death by rejuvenation, such as the immortal hydrozoan *Turritopsis nutricula*^1^, or may reveal prolonged life spans such as the Greenland shark, *Somniosus microcephalus*^2^, the black coral *Leiopathes glaberrima*^3^ and the sponge class *Hexactinellida*^4^. In fact, most multicellular organisms age through a single and directed senescence phenomenon in ontogeny that culminates in their inevitable deaths.

For aging, we studied here the model species *Botryllus schlosseri* (Fig. 1a,a’), a cosmopolitan marine colonial tunicate^5^, that has been emerged as an important species in diverse biological disciplines, including aging, allorecognition, and regeneration^6–9^. In addition to whole organismal aging, *Botryllus* presents well-studied rhythmic (weekly) cycles of life-and-death of its functioning modules (the zooids), each termed as a blastogenic cycle^10,11^ (Fig. 1b), a phenomenon considered as the ‘underwater phoenix’^12^. In each colony, three generations of modules co-exist, all genetically identical and asexually borne, altogether embedded within a gelatinous supporting matrix, the tunic (Fig. 1a’). The tunic further holds a ramified blood system connecting all modules to each other’s. The oldest generation of modules are the functioning zooids (Fig. 1a,b), that feed and breed. At the same time, the two younger generations, the first and secondary buds, experience fast ontogenic development (Fig. 1a,b). As in the Phoenix, the ancient Egyptian mythical bird that ignites itself to ashes and then emerges from its own ashes utterly renewed in endless cycles, each generation of zooids in *Botryllus* colonies are consumed by a weekly apoptotic wave^10^ and replaced by the swiftly matured generation of the first buds that becomes the next generation of functional zooids. Thus, *B. schlosseri* represents a colonial organism with repeated mosaic-like aging at the modules level and with a high regenerative power expressed by the replacement of modules that do not age according to classical fashion^13^. Aging at the whole-genet level (the sum of all ramets) may also occurs, and when develops it reflects a sharp contrast to the common pathways of senescence in unitary organisms^14–16^.

**Figure 1.**
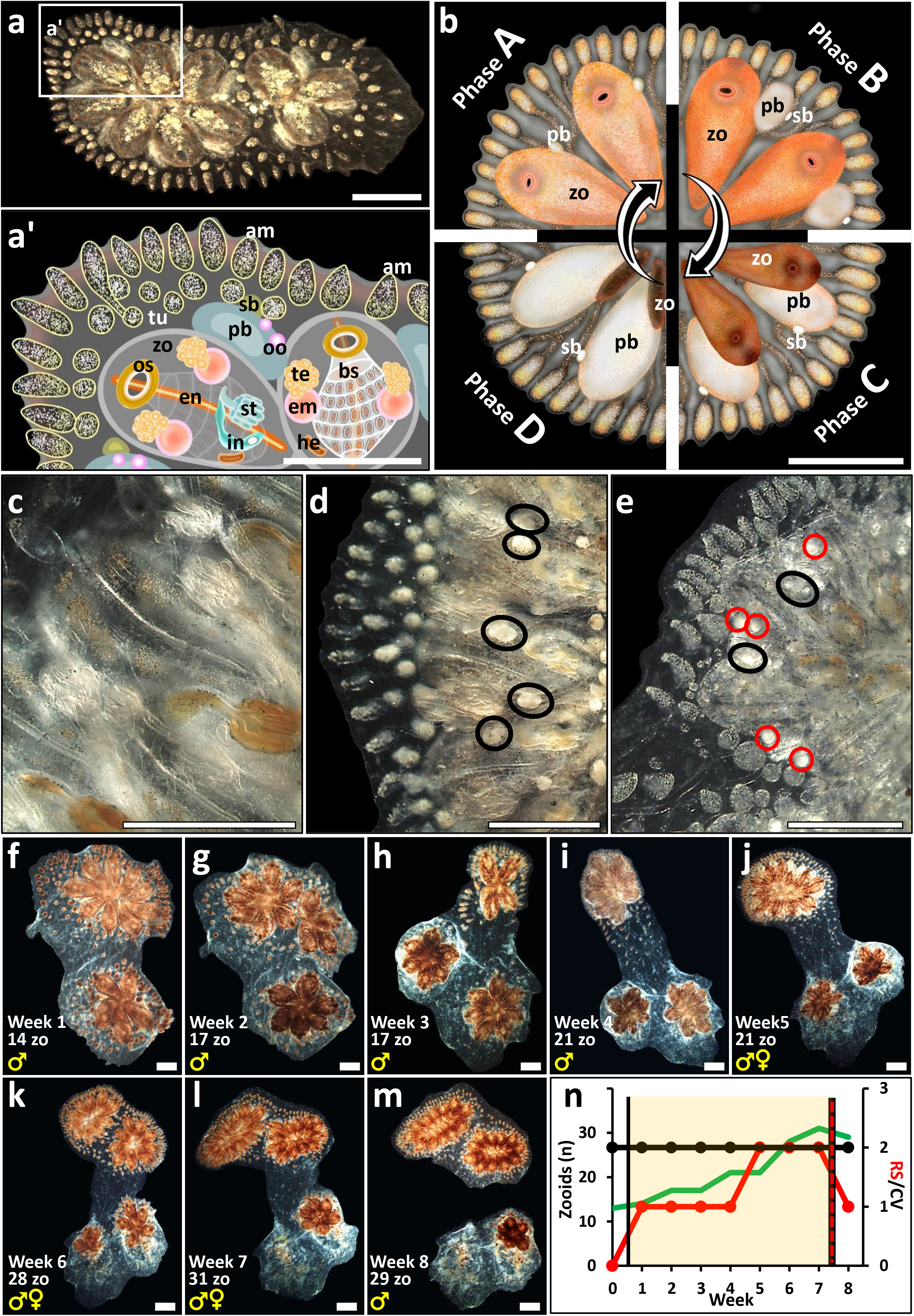
*Botryllus schlosseri* morphology and anatomy (a-e). (a) Mature colony composed of two flower-like shaped colonial systems, including 11 zooids. (a’) An illustrated magnification of Fig. 1a framed section, depicting the *Botryllus* colony organs. Am=ampullae; bs=branchial sac; em=embryo; en=endostyle; he=heart; in=intestine; os=oral siphon; oo=oocyte; pb=primary bud; st=stomach; sb=secondary bud; tu=tunic; zo=zooid. (b) Blastogenic cycle, a weekly recurring phenomenon of life- and-death of modules, divided into four phases, A to D, *sensu*^10,42^. The cycle encompasses three co-existing generations of modules: zooids, primary buds and secondary buds. Along blastogenesis, primary and secondary buds grow, but their oral siphons are not open yet as in functional zooids. At the end of each cycle (phase D or ‘takeover’) zooids are absorbed by an apoptotic wave and phagocytosis. The beginning of a new phase A is marked by the opening of the siphons of the primary set of buds. zo=zooid, pb=primary bud, sb=secondary bud. (c-e). Mature zooids presenting different reproductive statuses (RS). (c) Sterile zooid. (d) Male-only zooid. (e) Hermaphrodite zooid. Testes are black encircled; Oocytes are red encircled. (f-m). A colony along a single ‘astogenic segment’ (an *Orshina* segment) through 8 consecutive weekly observations (f-m). During the first four weeks (f-i), the colony is male-only (RS=1; red curve). and hermaphrodite (RS=2) along the next three weeks (j-l). In the eighth week observed, the colony return to a male only status (RS=1). In this example a single fission event was observed. (n) A schematic illustration assembling the three colonial demographic parameters documented in observations f-m. Borders (black vertical lines) are placed according to borders rules for an astogenic segment (yellow transparent square). X axis signifies the documented period (weeks). The left Y axis is for the zooid numbers (green curve) and the right Y axis for either the RS (red curve) or CV (black curve). Scale bars=1mm.

*B. schlosseri* colonies are further known to present two major life-history strategies, semelparity and iteroparity, each signified by co-evolved sets of trade-offs^17^. A recent study that followed *B. schlosseri* colonies from birth to death under relaxed laboratory conditions^18^ has elucidated three additional, yet distinct, life-history strategies, each disparate from the others, based on the existence/absence of colonial fission into two or more independent living ramets and fission properties: NF (no fission along the life span), FA (fission develops after the colony reaches its maximal number of zooids), and FB (fission develops before the colony reaches its maximal number of zooids (Fig. 2a-c).

**Figure 2.**
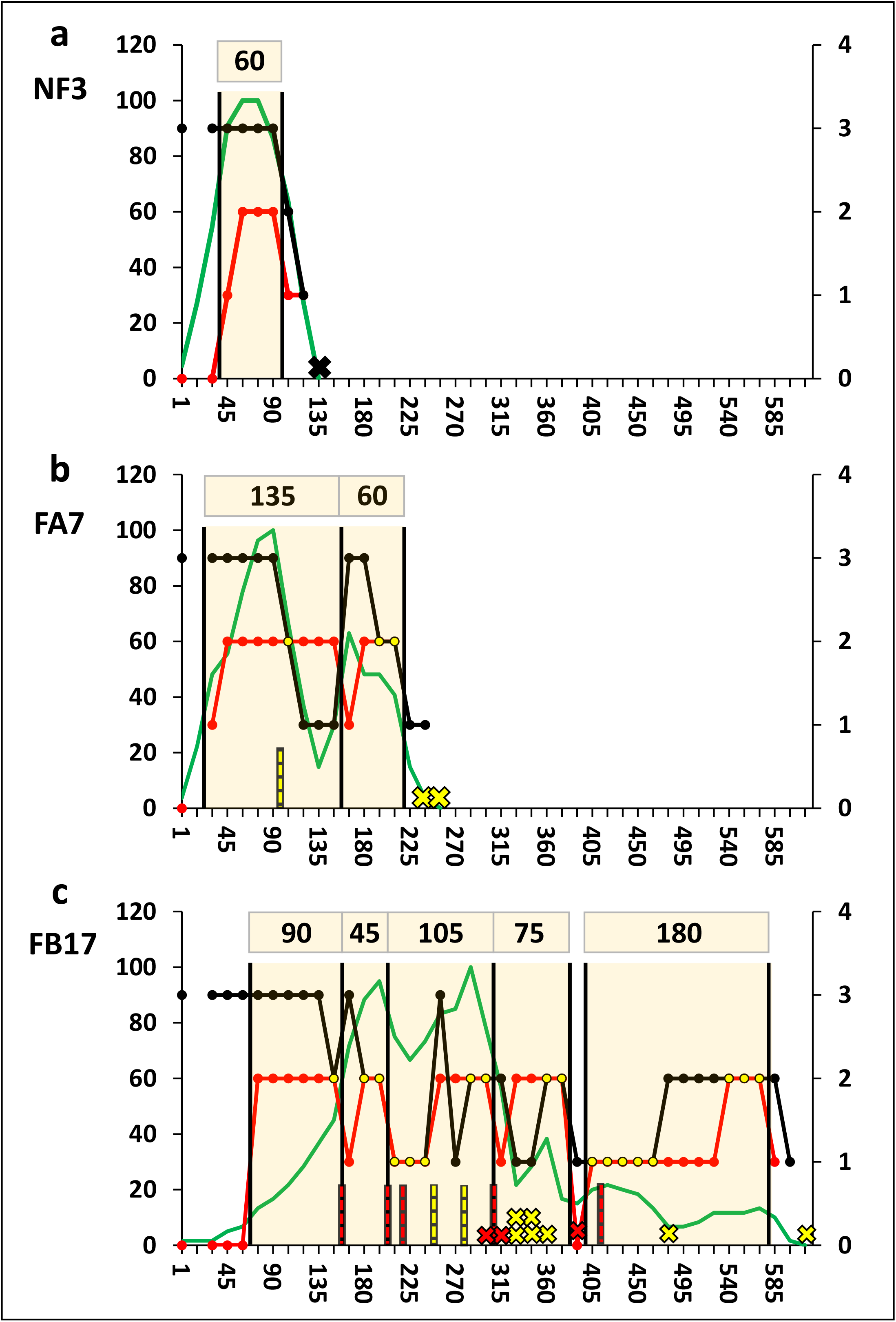
(a-c). Representative colonies that depict the ‘astogenic segments’ (yellow transparent squares) and the number of zooids, RS and CV for each one of the three life history strategies, NF, FA and FB. X axis signifies the documented period (days), the left Y axis is for the zooid numbers (green curve) and the right Y axis for either the RS (red curve) or CV (black curve). (a) A non fissioned type (NF), colony NF3. The colony presents a bell-shaped growth pattern, survived for a single RS cycle and started with maximal CV. (b) An FA type that fission after reaching peak size, colony FA7. The colony presents two bell-shaped growth patterns, each in a different ‘astogenic segment’, survived for two RS segments, and went through a single fission event at age of 3 months (divided into two ramets). The genet started with maximal CV, followed by a minimal CV period and rejuvenation to a second ‘astogenic segment’ that completed by the death of the two ramets in two consecutive observations. (c) An FB type that fission before reaching peak size, colony FB17. The colony presents three-four bell-shaped growth patterns, each in a different ‘astogenic segment’, survived for five RS segments, and went through seven fission events (divided into ten ramets). The colony started with maximal CV, and experienced 5 or 6 rejuvenation events along life. Death events are marked with crosses: red crosses - deaths located adjacent to the ‘astogenic segment’ borders, yellow crosses - deaths away from the ‘astogenic segment’ borders, black cross - a dead colony in NF. Fission events are marked with vertical dashed lines: red dashed lines - fissions adjacent to ‘astogenic segment’ borders, yellow dashed lines - fissions away from the ‘astogenic segment’ borders.

Studying in depth various demographic traits along the life spans of *Botryllus* colonies, we unveiled here a second phoenix-like phenomenon, developing on the colonial level. We noticed that sexual reproductive statuses (hermaphroditism, male only and sterile settings), colonial vigorousness and colonial sizes, altogether represent coinciding and repeated rhythms of astogenic segments on the whole-genet level. Each of the astogenic segments is termed as *Orshina*, symbolizing the immortal Jewish mystical bird^19^ and the sum of all segments as the *Orshina* rhythm. The *Orshina* rhythm is thus typified by scheduled orchestrated morphological phenomena of colonial degradation, rejuvenation, reproduction, fission and death of ramets.

## Materials and methods

### Animals

Colonies of *Botryllus schlosseri* were reared and kept under constant temperature (20°C) and light/dark regimen (12:12 hours) in 16L standing seawater plastic tanks, at the National Institute of Oceanography (Haifa, Israel) facility, as described^20^. Colonies were followed from birth to death and observations were performed every 15±5 days as from age of 1 month, under Nikon stereo microscope (SMZ 1000).

#### Documented parameters included

(a) Number of zooids: Active zooids were counted each observation for each genet, e.g., following fissions events, the total number of zooids in all ramets were combined; (b) Colonial Vigorousness (CV): A semi-qualitative score, averaging the morphological statuses (ranging 1-3) for each one of the three major colonial compartments, the zooids/buds, the peripheral ampullae and the tunic. CV scores ranged between 1 to 3, where 3 signifies best vigorousness, 2 is an intermediate status and 1 is the lowest (*sensu*^18^ Suppl. Fig. 1); (c) Reproductive Status (RS): Colonies of *B. schlosseri* present alternate sexual status (sterile, male only, hermaphrodites) along life span. At onset, colonies are sterile, then male gonads appear first along ontogeny, and female gonads appear in the following 2-4 blastogenic cycles (Fig. 1c-e). This hermaphroditic phase is followed by periods of sexual sterility and/or only male status^20,21^. The RS is a genotype specific status which is varied between different colonies under the same environmental settings^21^. RS is a semi-quantitative score, where grade 0 is for ‘sexual sterility’ status, grade 1 for ‘only male’ colonies, and grade 2 for ‘hermaphrodite status’, irrespective to the number of female/male gonads per zooid/bud; (d) Colony fission: Colonial fission (Fig. 1f-n) is an astogenic process ending in the splitting of a colony into two or more ramets, a well-documented process in *B. schlosseri* colonies^21^. As fission gradually develops (few days to weeks from commence), the exact date of fission is determined as from the first documented cut-off of blood vessels between splitting ramets, even though they may still be physically connected by degraded tunic matrix.

### Statistical analyses

Spearman’s correlation coefficient was calculated between the three documented traits (zooid numbers, CV and RS) for each colony. Then the average and standard deviation of Spearman correlation coefficients of all colonies of the same type of life history strategy (NF, FA, FB) were calculated. One sample T test was used to examine if each average correlation between each couple of parameters for each life history type was significantly different from zero. On fissioned colonies, Chi square tests were designed to evaluate: (a) Non-random death of ramets, performed on simple cases of fission events. (b) Deaths of ramets, associations with astogenic parameters. (c) Fission events, associations with astogenic borders. Chi-square tests analyses were performed using Microsoft Excel program. Examining the possibility that deaths events are dependent samples, Wilcoxon sign test was formed using SPSS program and was designed for an additional evaluation of ramets death closeness to astogenic borders.

## Results

### General

The documented parameters, number of zooids, RS and CV for each of the 81 colonies (NF=35; FA=23; FB=23) were organized individually (examples in Fig. 2a-c; All graphs in Suppl. Figs. 1-3). Focusing first on RS, we noticed repeated cycles of reproductive statuses along the colonies’ life spans where the first wave starts with a sterile period. A male-only status emerges later, trailed by a period of hermaphroditic status. The length of male-only period to hermaphrodite period is regarded as a single segment. The sterile period at the beginning of life was precluded from segments delineation, since was documented in only 8 short term midlife periods (in colonies NF30, NF32, FA22, FB2, FB5, FB10, FB17, FB20; Suppl. Figs. 1-3). Each of the RS cycles (called hereby astogenic segment) was delineated (vertical lines in Fig. 2a-c). Each astogenic segment length averaged 89±45 days (n=142 segments in 70 colonies in which clear segments were established), and examples representing the NF, FA and FB life history strategies are depicted in Fig. 2a-c. Gradual differences between segment lengths among the life history strategies, albeit non-significant, were documented, where NF showed the shortest length (75±31, n=30 segments), FA an intermediate length (86±38, n=42 segments), and FB presented the longest segment length (98±53, n=70 segments). Interestingly, hermaphroditism spanned about 2/3 of the astogenic segment length in all, NF, FA and FB colonies. Interestingly, the three types presented constant and almost fixed RS ratio (Fig. 3; Suppl. Table 2).

**Figure 3.**
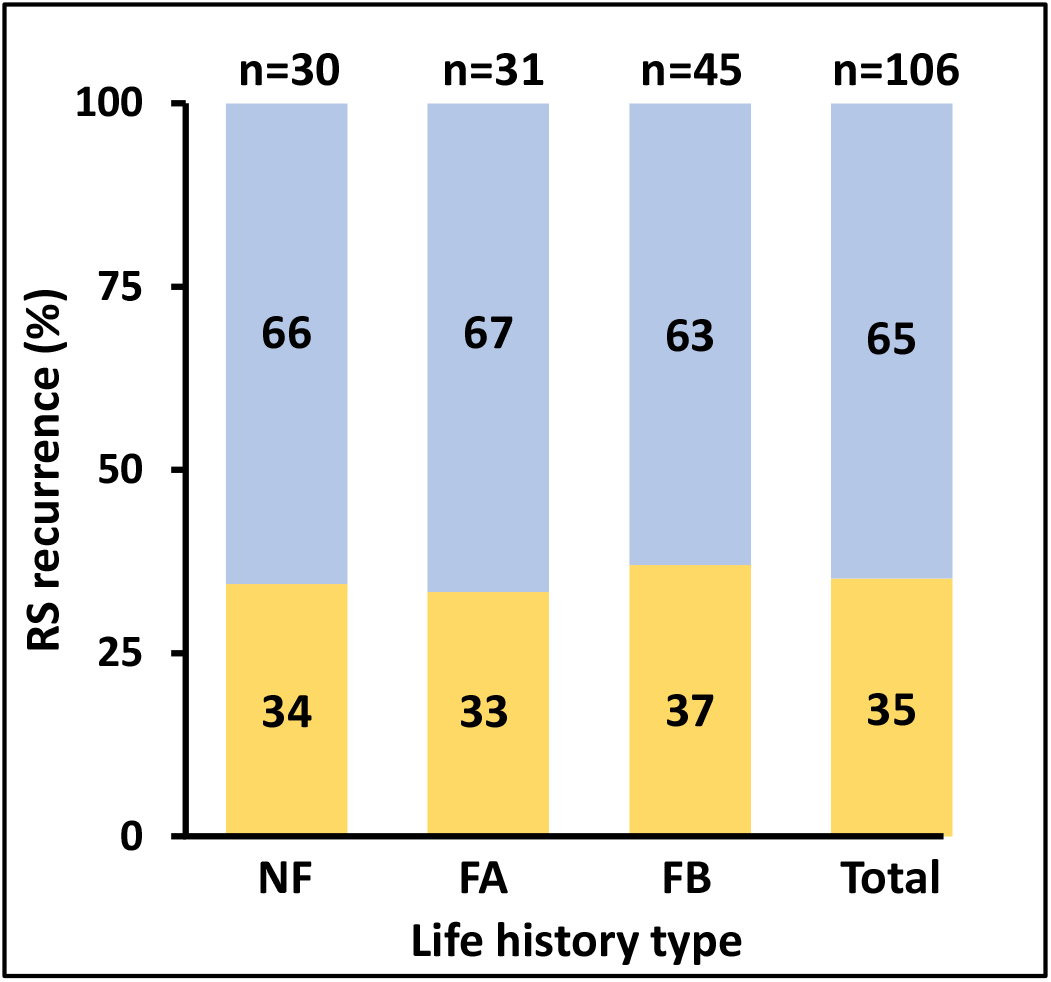
Male only status (yellow) and hermaphroditic status (blue) distributions (%) in typical ‘astogenic segments’ of the three life history types (NF, FA and FB) and the average summary (total) for *B. Schlosseri* colonies. The number of total segments studied for each life history type are shown above the columns.

A high correlation was found between the number of astogenic segments and life span lengths (Pearson correlation coefficient 0.8; n=77; *p*<0.05). NF strategy is crowned by 1.3±0.6 astogenic segments (n=32) along the colonies’ life spans, followed by FA strategy (2.2±1.5 segments; n=22) and FB strategy (3.2±1.3 segments; n=23). Pairwise Spearman’s correlation coefficient was calculated for each pair of parameters along the life of each colony, summing up to 50 observations per pair (no. of zooids vs. RS; no. of zooids vs. CV; RS vs. CV; Suppl. Table 1a), followed by One-sample T test analyses to test the significance of the average correlations for each pair of parameters in each life history strategy. The results revealed significant correlations, mostly positive, for most pairs (Suppl. Table 1b). While in NF all Zooids-RS, Zooids-CV and RS-CV correlations were significant (*p*<0.005), in FA and FB, two pairs were significant, Zooids-RS, Zooids-CV, and Zooids-RS, RS-CV, respectively (Suppl. Table 1b). In the three life strategies, zooids vs. RS and zooids vs. CV showed positive correlations, while the pair RS vs. CV showed a negative correlation, suggesting antagonist expressions (Suppl. Table 1a). In the four longest-living colonies FB 20-23 (life spans of 623, 458, 471 and 726 days, respectively) the three pairwise combination tests were significant (*p*<0.05; apart from the pair ‘zooids vs. CV’ in colony FB21).

### Ramet’s death and fission are synchronized with the astogenic segments

We then focused on fission events and ramets’ deaths. FA and FB colonies went through fission events, forming naturally detached ramets that survived for few days to prolonged periods (deaths are marked in crosses in Fig. 2a-c), altogether constructed either simple cases of a single fission event into just two ramets (Fig. 4a) or complex cases of multiple ramets per genet (Fig. 4b). For statistical analyses on ramets’ life spans we focused only on colonies that fissioned just once (n=14 cases; FA and FB types; Suppl. Table 3). Of these, 50% of the coupled ramets died simultaneously, non-randomly, meaning that both ramets/genet died together (documented dead in the same or in a consecutive observation). Statistical analyses showed significant likelihood of the ramets to die at the same time (Chi square test, *p*<0.001, n=14, df=1; Suppl. Table 4).

**Figure 4.**
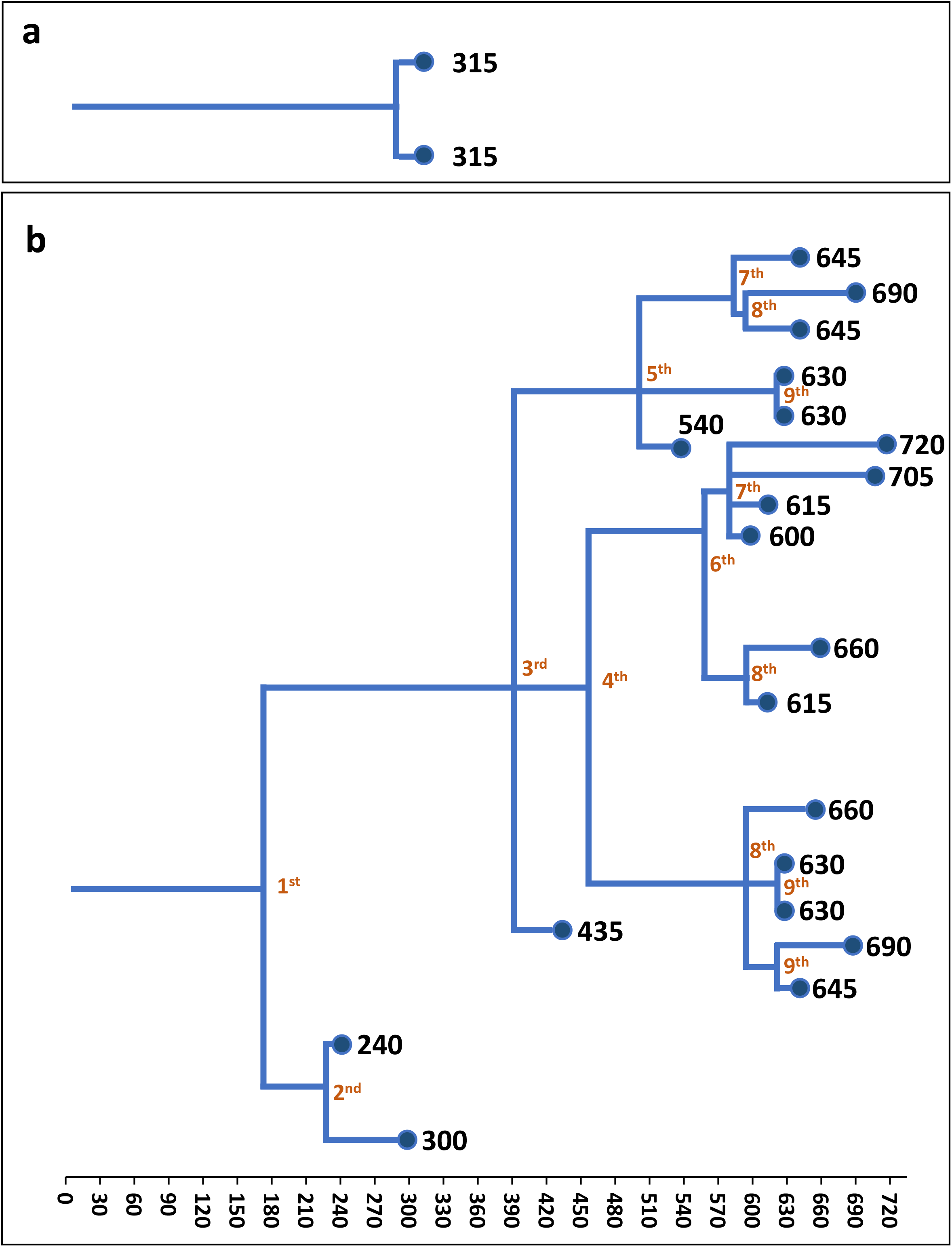
Fission sequences in *B. Schlosseri* colonies. (a) A colony that fissioned only once (FA16). (b) A colony (colony FB3) that went through 9 fission events, creating 20 ramets. Black numbers depict life-long periods (days) for each ramet. Orange numbers show chronological sequence of the fission events.

Observations further revealed that ramet’ deaths were aligned, in many cases, with the astogenic segments’ borders. An example with two ramets that died adjacent to the segment borders is depicted (Fig. 2c; red crosses). As colonies were observed every 15 days and as borders are delineated between observations, a death associated with the segment border was assigned to ramets dying up to 7.5 days before or after a border line. Since the segment borders cannot be delineated at the end of genets’ life spans, we did not include death cases allied with the genet demise in the Chi square analyses, thus included death cases occurring up to the middle length of the last astogenic segment (45 days). Results revealed that ramets deaths were significantly adjacent to the astogenic borders (Chi square, *p*<0.0001, df=1, n=126 ramets deaths in 29 genotypes with approved segments; Suppl. Table 5). Wilcoxon sign test further confirmed non-random increased deaths at the segment borders (*p*<0.0001, z=-3.49, n=126 ramets deaths in 29 genets), signifying that ramet’s deaths are matched with segments outskirts.

Observations further revealed many cases where fission events were aligned with the segments’ borders. An example with five fission events that occurred adjacent to the segment borders is depicted (Fig. 2c; dashed red lines). As both borders and fissions are delineated between observations, a fission event associated with a border was assigned to ramets dying up to 15 days before or after the border line. As above, we included fission events occurring up to the middle length of the last astogenic segment (45 days). Results revealed that fission events were significantly adjacent to the astogenic borders (Chi square test, *p*<0.0001, df=1, n=165 fissions in 41 genets; Suppl. Table 6). Wilcoxon sign test did not discern significant relationship (*p*>0.05) between the astogenic segment outskirts and fission events.

### The astogenic rhythm

To assess the rhythmic nature of astogenic segments along the whole life spans of the colonies, we developed a method that enables the transformation of the observed growth rates (changes in the number of zooids between two observations) into ‘growth trends’ (measured as positive [+] or negative [-] change in zooid numbers between two observations). This allows the comparison of growth between colonies, regardless the number of zooids. For this purpose, the data presented in the individual growth charts (Suppl. Fig. 1-3) was modified as followed: A numeral ‘1’ was assigned to the first observation of the first settled zooid. We then added the value ‘1’ any time the number of zooids at the following observation was higher than in the former observation (e.g., 2), and reduced the value ‘1’ from the observation value where the next census revealed lower number of zooids (e.g., 0). Growth trends curves for the 81 genets (Fig. 5; green curves) were further aligned with RS and CV curves and divided into three groups: 1. Colonies exhibiting life span of a single astogenic segment; 2. Colonies with two astogenic segments; 3. Colonies with three or more astogenic segments along their lives. For each of the 81 genotypes, the three life history types are further highlighted (NF-yellow frames; FA-blue frames; FB-red frames; Fig. 5).

**Figure 5.**
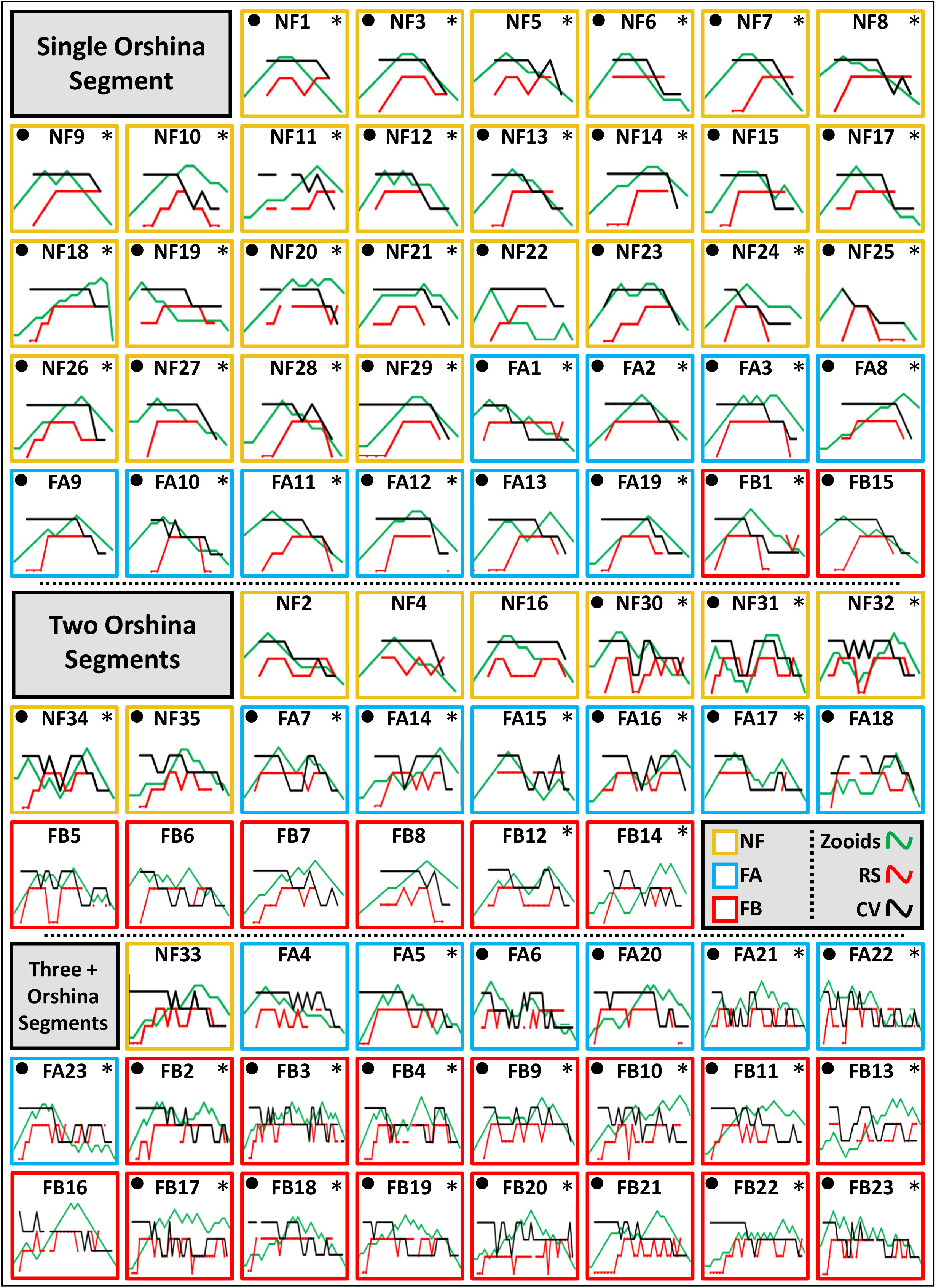
All studied genotypes (n=81) that were followed from birth to death in accordance with the three life history types (NF =yellow frames; FA =blue frames; FB =red frames). Three demographic features are depicted for each genotype: growth trends of zooids (green curves), RS (red curves) and CV (black curves). The genotypes are divided into three groups: 1. Colonies with a single astogenic segment 2. Colonies with two astogenic segments; 3. Colonies with three or more astogenic segments. Genotypes where the number of segments matched number of zooids peaks are marked with asterisks. genotypes where the number of segments matched with the number of CV peaks are marked with black circles.

Results (Fig. 5) reveal that NF colonies are more prominent in the ‘single astogenic segment’ group (60% of all genotypes), FA colonies are divided more or less equally between the three astogenic groups and FB colonies are more evident in the third astogenic group (70% of all genotypes). Using the ‘growth trends’ analyses in the group ‘single astogenic segment’, 84% of colonies further show a single peak of zooids (32/38 colonies) compared to 55% of colonies presenting two clear peaks (11/20 colonies) in the ‘two astogenic segments’ group, and 74% of colonies presenting several peaks (17/23 colonies) in the ‘3+ astogenic segments’ group (marked with asterisks in Fig. 5).

Analyses were then performed on Colonial Vigorousness (CV) peaks along the life span of each genotype. About 84% (32/38) of ‘single astogenic segment’ colonies presented a single CV peak, 45% (9/20) of the ‘two astogenic segments’ colonies showed two CV peaks and 83% (19/23) of ‘3+ astogenic segments’ colonies present several CV peaks (marked by black circles; Fig. 5). Remarkably, 60% (49/81) of the colonies showed a match between the numbers of astogenic segments, the number of zooidal peaks and the number of CV peaks, an indication for a rhythmic nature along the colonies’ life spans (marked both in asterisks and circles; Fig. 5).

### The four stages of the astogenic rhythm

The analyses of all astogenic segments in all 81 colonies studied (Fig. 2; Fig. 5; Suppl. Fig. 1-3), revealed common morphometric features to all three life history types (NF, FA, FB), in relation to the number of zooids, RS and CV, that are characterized by repeated astogenic segments, each divided into four developmental phases, α, β, γ and δ (Fig. 6). In addition, during the first 35% of the length of each segment, the colony is ‘male only’, and then, for the rest 65% of the segment, it reveals the hermaphroditic state (Fig. 3; Suppl. Table 2; the first segment is delineated as from the end of the juvenile sterile phase). In the astogenic segment phase α, the colony has a high CV status, presenting male only gonads. The astogenic segment phase β is a developmental state where the colony is hermaphrodite with higher CV status and represents a peak number of zooids. In astogenic segment phase γ, the colony typically presents a reduced CV status, hermaphroditic state and gradually reduced number of zooids. In the fissioned types this is also the phase where colonial fission develops. Phase δ is typified with gradually reduced CV status, reduced hermaphroditism and also with colonial fission events (when developed). The end of phase δ is the ‘turn point’ or the pivotal phase where the ramet/genet either die (most NF colonies, ramets of FA/FB genotypes, or entire FA/FB genotypes) or rejuvenate into phase α of the next astogenic segment (ramets and/or whole FA/FB genotypes; (Fig. 6). Having these results (Fig. 6), each of the astogenic segments in the life span of a colony is called an ‘*Orshina* segment’ and the sum of repeating astogenic segments per a *Botryllus* genotype, is termed as the ‘*Orshina* rhythm’.

**Figure 6.**
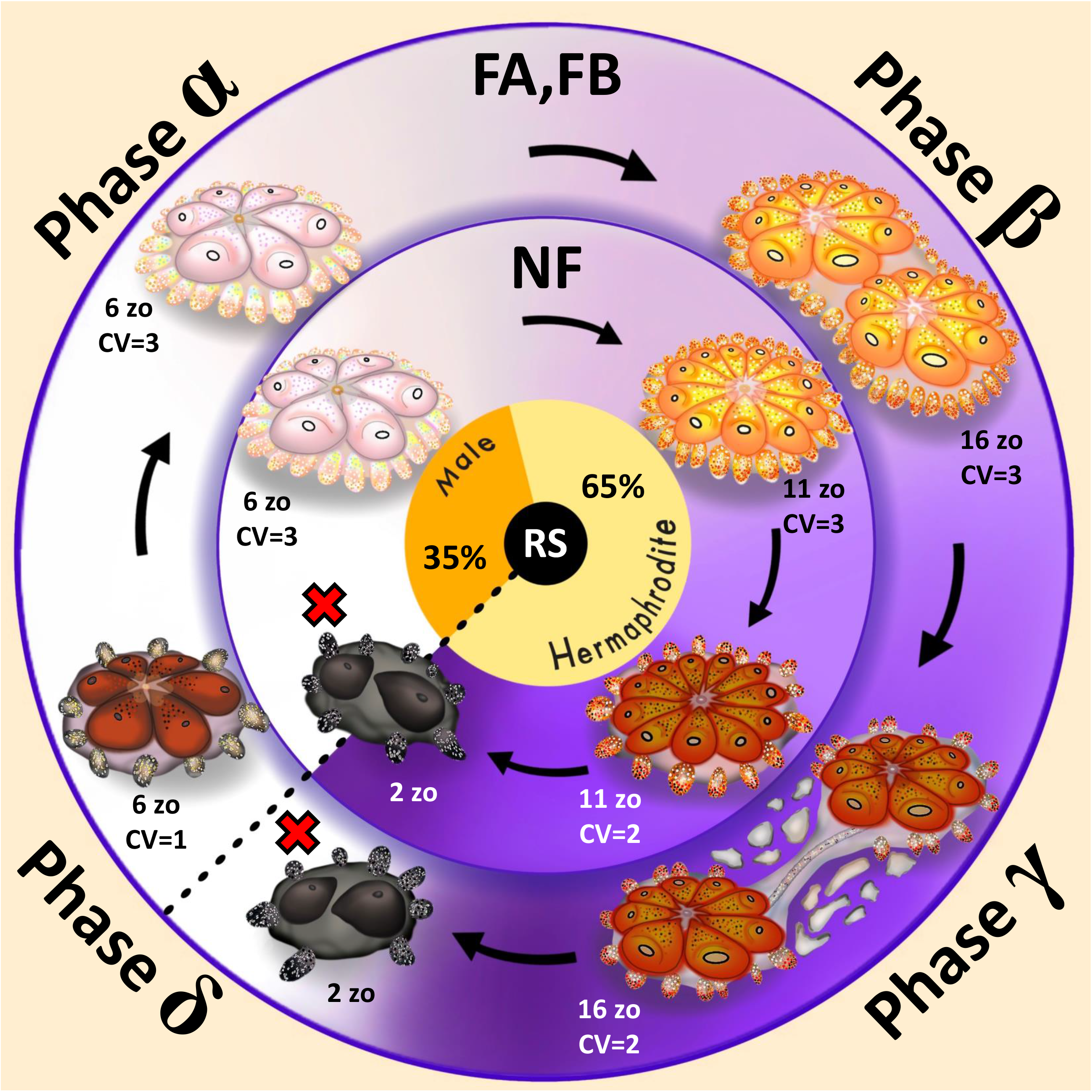
Summary of *Orshina* segment/s in non-fissioned colonies (NF, inner cycle) and in fissioned colonies (FA, FB, outer cycle). Each *Orshina* segment is divided into four phases, α through δ. In NF colonies, the rhythm repeats primarily only once, and in the fissioned colonies the rhythm usually repeats for two to three times (under lab conditions) until final death of the last ramet. The following description of the phases concern both inner and outer cycles. The reproductive status, RS, is shown in the innermost cycle, relating to the statuses along all phases. Phase α: the colony presents male-only gonads and high CV status (CV=3). Phase β: the developmental phase where colony’s size is greater than in phase α. Hermaphrodite status and high CV statuses are common. Phase γ: while hermaphrodite status is present, CV status may be lower than in phase β. In the fissioned types the tunic between the systems is gradually deteriorating. Phase δ: NF colony usually dies at the end of this phase. Both death of ramets and fission events appear in this phase. Survived ramets present smaller size, lower CV, and then they go through rejuvenation at the end of this phase, enabling the ramets to carry on a new *Orshina* segment. The *Orshina* border is marked by a dashed line. Numbers of zooids presented are a suggested simplification. Different colonies present different sizes, shapes and colors. Red crosses=dead entities; zo=zooids.

## Discussion

The name *Orshina* is taken from the esoteric Jewish mysticism essays (in the Babylonian Talmud), referring to a bird that was secured in Noah’s arch. According to the legend, *Orshina* remained silent while Noah supplied food for the rest of the survived animals, and for its politeness, *Orshina* was blessed with immortal life, where every one-thousand years it degrades and then rejuvenates^19^. As a metaphor, we tagged the genet level’s life and death cycles in *Botryllus* colonies as the *Orshina* rhythm and each cycle as an *Orshina* segment (Fig. 6). Each such segment lasts ca. 3 month long, is not linked to the weekly life and death blastogenic cycles developing at the modules level, and exists in all three *Botryllus schlosseri* life history strategies (NF - colonies that never fissioned; FA – colonies that fissioned after maximal peak of zooids; FB - colonies that fissioned before the maximal peak of zooids^18^). Here we further elucidated that the *Orshina* rhythm is orchestrated by three traits at the colony level (expressed in different ramets and/or the whole genet), the colonial reproduction status (RS), the genet’s size (total number of zooids) and the colonial vigorousness status (CV). Each of these colonial traits portrays repeating archetypal curves over time, and all traits are further synchronized with each other, making an overall rhythm of reproduction, colonial fission, life/death, and aging/rejuvenation phenomena.

As specified, an *Orshina* rhythm is composed of one or more *Orshina* segments (each 89±45 days long), each is represented by four development phases (α, β, γ and δ; Fig. 6). Regardless of the life history strategy, or the considerable variance of segments lengths, the male and the hermaphrodite phases within an *Orshina* segment occupy 35% and 65% of the *Orshina* segment length, respectively (Figs. 3,6), implying uniformity of reproductive statuses. An *Orshina* segment is also typified with a characteristic bell-shaped colonial size, where maximal size occurs at the middle of a segment.

Any *B. schlosseri* genet may exhibit several constructed aging/rejuvenation phenomena (some of them simultaneously occurring), such as semelparity (e.g. single reproductive episode) versus iteroparity (e.g. multiple reproductive sequels)^17^, programmed life span as compared to wear-and-tear aging processes^14,16^, weekly aging of colonial modules (blastogenesis)^22^, rejuvenation^23–26^, and the immortality of germ/somatic cell lines^27^. The above phenomena reflect the existence of spatial and seemingly stochastic age-mosaic modules within a genotype, the regulation of aging by an extreme regeneration power, and the replacement of somatic modules (*sensu* the disposable soma tenet^13,28^), all targeting a modular, colonial species that does not age according to the common aging phenomena in unitary organisms^14–16^.

Considering the *disposable soma theory*, aging is considered as the inevitable outcome of decisions performed along the organism life span, all associated with the distribution of energy sources between the soma and the germ line. As long as the organism is sexually sterile, sources are allocated to growth, maintenance, repair, storage and defense; whereas sexual reproduction diverges the sources towards the germline, further imposing senescence^29–32^. While the above holds for unitary organisms that display germline sequestering, in colonial organisms that do not sequester their germline (such as *Botryllus*^33^), evolutionary trends may shift to biological statuses where the soma is not carefully maintained nor efficiently repaired (e.g.,^24,28,34^), somatic constituents that are replaced on a regular basis, and biological statuses that exhibit distinctive senescence occurrences^13^.

In *Botryllus*, most NF colonies present a single *Orshina* segment along their lives (Fig. 6, inner circle) and then they die. The fissioned colonies in FA and FB life history strategies (Fig. 6, outer circle) hold two or more *Orshina* segments, where at the end of each segment, rejuvenation start off, leading to a new *Orshina* segment. The pivotal rejuvenation points (at *Orshina* phase δ) represent a clear shift from colonial deterioration to growth and development that emerges following the completion of colonial fission events, coinciding with the death of ramets that fail to transcend this critical turn-point in astogeny. Thus, ‘fission’ at the genet level may have bearing on longevity. This resembles mechanically fissioned annelids where repeated fissions extended life spans^35^, fission augmented growth in ramets of didemnid ascidians^36^, and elongated telomeres in the fissiparous starfish *Coscinasterias*^37^.

The literature in botryllid ascidians thoroughly discusses the astogenic phenomenon of blastogenesis (Fig. 1b), the weekly recurrent life and death events developing at the modules level^11,12,27,38–41^. Basically, the *Orshina* segment described here, is a newly elucidated life and death event developing at the genet level, lasts about three months on the average (containing ca. 13 blastogenic cycles), ends by either the death of the whole colony or by rejuvenation, and manipulated by absence/existing fission events on the colony level, all revealing comparable and disparate properties when compared to blastogenesis (Table 1). A blastogenic cycle, in turn, operates on the zooids level, lasts about a single week, ends with the complete eradication of existing functional zooids and their replacement by a new developing set of primary buds (the take-over phase) and fission occurs at the zooidal-system level (Table 1). Yet, the two phenomena, blastogenesis and *Orshina* rhythm seem to be “programed” by similar coding rules, *sensu* “As above so below, as below, so above” (*Hermes Trismegistus* in “Hermetic Corpus,”).

**Table 1.**
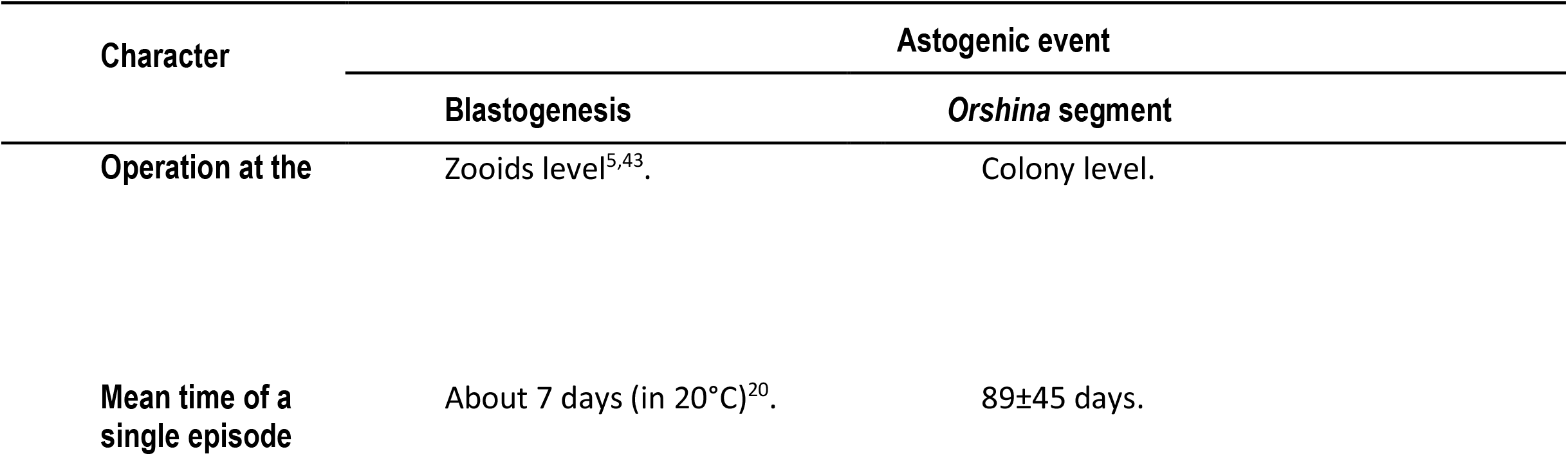
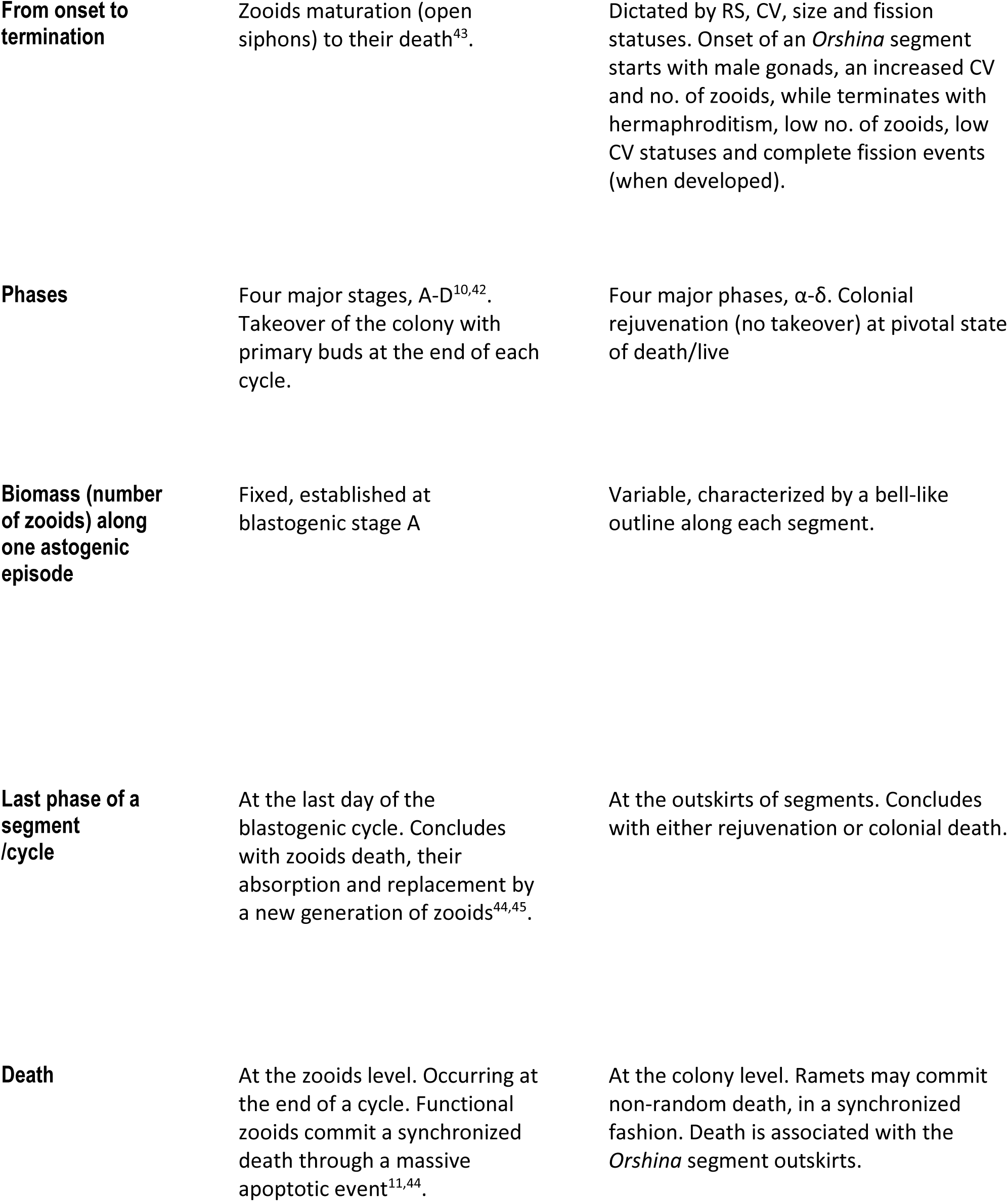
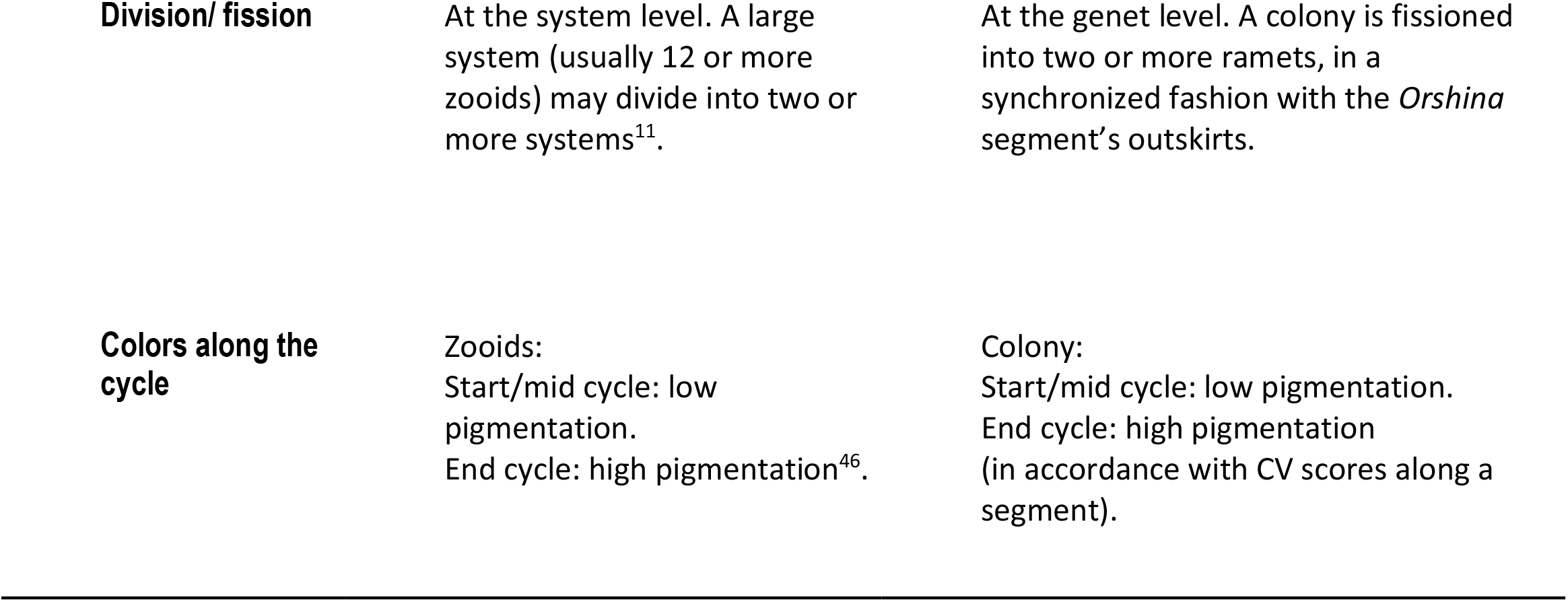
Comparative attributes for a blastogenic cycle and the *Orshina* segment.

The average life span of *Botryllus* colonies in our experiments (290 days^18^) composed of 41 blastogenic cycles and 2.1 *Orshina* segments. *Botryllus schlosseri* genets thrive through rhythmic astogenic (*Orshina*) segments, each roughly lasts for three months that are most likely programmed by inherent and conserved genetics. Given optimal environmental conditions, a *Botryllus* colony, like *Orshina* the bird, is potentially able to live for long periods and rejuvenate in a rhythmic manner, a notion supported by the literature, where genets live for >20 years^14^ or die through a programmed life span^16^. The current findings indicate that reproduction, life span, death, rejuvenation and fission events are, at least in part, scheduled processes along the constructed *Orshina* rhythm.

## Acknowledgement

We thank G. Paz for support along the way. O.B.H thanks her parents for financial support.

## Fund support

BR was supported by a grant from the United States-Israel Binational Science Foundation (BSF no. 2015012) and a grant from the Israel Science Foundation (ISF No. 172/17).

## Author contribution

Conceptualization and designing of the experiments and the ideas by O.B.H., R.B.S. and B.R.. O.B.H. and B.R. wrote the main manuscript. I.I. supervised the statistical analyses; all authors edited the manuscript.

## References

1. Bavestrello, G., Sommer, C. & Sarà, M. Bi-directional conversion in Turritopsis nutricula (Hydrozoa). Sci. Mar. 56, 137–140 (1992).

2. Nielsen, J. et al. Eye lens radiocarbon reveals centuries of longevity in the Greenland shark (Somniosus microcephalus). Science 353, 702–704 (2016).

3. Roark, E. B., Guilderson, T. P., Dunbar, R. B., Fallon, S. J. & Mucciarone, D. A. Extreme longevity in proteinaceous deep-sea corals. Proc. Natl. Acad. Sci. 106, 5204–5208 (2009).

4. Conway, K. W., Barrie, J. V., Austin, W. C. & Luternauer, J. L. Holocene sponge bioherms on the western Canadian continental shelf. Cont. Shelf Res. 11, 771–790 (1991).

5. Ben-Hamo, O. & Rinkevich, B. Botryllus schlosseri—A model colonial species in basic and applied studies. in Handbook of Marine Model Organisms in Experimental Biology 385–402 (CRC Press, 2021).

6. Blanchoud, S., Rinkevich, B. & Wilson, M. J. Whole-Body Regeneration in the Colonial Tunicate Botrylloides leachii. in Marine Organisms as Model Systems in Biology and Medicine (eds. Kloc, M. & Kubiak, J. Z.) vol. 65 337–355 (Springer International Publishing, 2018).

7. Magor, B. G., Tomoso, A., Rinkevich, B. & Weissman, I. L. Allorecognition in colonial tunicates: protection against predatory cell lineages? Immunol. Rev. 167, 69–79 (1999).

8. Rinkevich, B. Primitive immune systems: Are your ways my ways? Immunol. Rev. 198, 25–35 (2004).

9. Rinkevich, B. Natural chimerism in colonial urochordates. J. Exp. Mar. Biol. Ecol. 322, 93–109 (2005).

10. Lauzon, R. J., Ishizuka, K. J. & Weissman, I. L. Cyclical Generation and Degeneration of Organs in a Colonial Urochordate Involves Crosstalk between Old and New: A Model for Development and Regeneration. Dev. Biol. 249, 333–348 (2002).

11. Manni, L. et al. Sixty years of experimental studies on the blastogenesis of the colonial tunicate Botryllus schlosseri. Dev. Biol. 448, 293–308 (2019).

12. Rinkevich, B. The tail of the underwater phoenix. Dev. Biol. 448, 291–292 (2019).

13. Rinkevich, B. 11 Senescence in Modular Animals. in The Evolution of Senescence in the Tree of Life (eds. Shefferson, R. P., Jones, O. R. & Salguero-Gómez, R.) (Cambridge University Press, 2017).

14. Anselmi, C. et al. Two distinct evolutionary conserved neural degeneration pathways characterized in a colonial chordate. Proc. Natl. Acad. Sci. 119, e2203032119 (2022).

15. Lauzon, R. J., Rinkevich, B., Patton, C. & Weissman, I. L. A morphological study of nonrandom senescence in a colonial urochordate. Biol. Bull. 198, 367–378 (2000).

16. Rinkevich, B., Lauzon, R. J., Brown, B. W. & Weissman, I. L. Evidence for a programmed life span in a colonial protochordate. Proc. Natl. Acad. Sci. 89, 3546–3550 (1992).

17. Grosberg, R. K. Life-history variation within a population of the colonial ascidian Botryllus schlosseri. 1. The genetic and environmental control of seasonal variation. Evolution 42, 900–920 (1988).

18. Ben-Hamo, O., Izhaki, I., Ben-Shlomo, R. & Rinkevich, B. Fission in a colonial marine invertebrate signifies unique life history strategies rather than being a demographic trait. Sci. Rep. 12, 15117 (2022).

19. Shemesh, A. O. Religious Literature, The realistic, and the Fantastic. Estud. Religiao 33, 235–255 (2019).

20. Rinkevich, B. & Shapira, M. An improved diet for inland broodstock and the establishment of an inbred line from Botryllus schlosseri, a colonial sea squirt (Ascidiacea). Aquat. Living Resour. 11, 163–171 (1998).

21. Rinkevich, B., Porat, R. & Goren, M. On the development and reproduction of Botryllus schlosseri (Tunicata) colonies from the eastern Mediterranean Sea: plasticity of life history traits. Invertebr. Reprod. Dev. 34, 207–218 (1998).

22. Ben-Hamo, O., Rosner, A., Rabinowitz, C., Oren, M. & Rinkevich, B. Coupling astogenic aging in the colonial tunicate Botryllus schlosseri with the stress protein mortalin. Dev. Biol. 433, 33–46 (2018).

23. Laird, D. J. & Weissman, I. L. Telomerase maintained in self-renewing tissues during serial regeneration of the urochordate Botryllus schlosseri. Dev. Biol. 273, 185–194 (2004).

24. Rinkevich, B. & Weissman, I. L. Botryllus schlosseri (tunicata) whole colony irradiation: Do senescent zooid resorption and immunological resorption involve similar recognition events? J. Exp. Zool. 253, 189–201 (1990).

25. Rosner, A., Kravchenko, O. & Rinkevich, B. IAP genes partake weighty roles in the astogeny and whole body regeneration in the colonial urochordate Botryllus schlosseri. Dev. Biol. 448, 320–341 (2019).

26. Voskoboynik, A., Reznick, A. Z. & Rinkevich, B. Rejuvenescence and extension of an urochordate life span following a single, acute administration of an anti-oxidant, butylated hydroxytoluene. Mech. Ageing Dev. 123, 1203–1210 (2002).

27. Rinkevich, Y. et al. Repeated, Long-Term Cycling of Putative Stem Cells between Niches in a Basal Chordate. Dev. Cell 24, 76–88 (2013).

28. Qarri, A., Rosner, A., Rabinowitz, C. & Rinkevich, B. UV-B radiation bearings on ephemeral soma in the shallow water tunicate Botryllus schlosseri. Ecotoxicol. Environ. Saf. 196, 110489 (2020).

29. Kirkwood, T. B. L. Evolution of ageing. Nature 270, 301–304 (1977).

30. Kirkwood, T. B. L. Evolution of ageing. Mech. Ageing Dev. 123, 737–745 (2002).

31. Kirkwood, T. B. L. & Holliday, R. The evolution of ageing and longevity. Proc. R. Soc. Lond. B Biol. Sci. 205, 531–546 (1979).

32. Westendorp, R. G. J. & Kirkwood, T. B. L. Human longevity at the cost of reproductive success. Nature 396, 743–746 (1998).

33. Rosner, A., Moiseeva, E., Rinkevich, Y., Lapidot, Z. & Rinkevich, B. Vasa and the germ line lineage in a colonial urochordate. Dev. Biol. 331, 113–128 (2009).

34. Svanfeldt, K., Lundqvist, L., Rabinowitz, C., Sköld, H. N. & Rinkevich, B. Repair of UV-induced DNA damage in shallow water colonial marine species. J. Exp. Mar. Biol. Ecol. 452, 40–46 (2014).

35. Martinez, D. E. Rejuvenation of the disposable soma: Repeated injury extends lifespan in an asexual annelid. Exp. Gerontol. 31, 699–704 (1996).

36. Ryland, J. S., Wigley, R. A. & Muirhead, A. Ecology and colonial dynamics of some Pacific reef flat Didemnidae (Ascidiacea). Zool. J. Linn. Soc. 80, 261–282 (1984).

37. Garcia-Cisneros, A. et al. Long telomeres are associated with clonality in wild populations of the fissiparous starfish Coscinasterias tenuispina. Heredity 115, 437–443 (2015).

38. Manni, L., Zaniolo, G., Cima, F., Burighel, P. & Ballarin, L. Botryllus schlosseri: A model ascidian for the study of asexual reproduction. Dev. Dyn. 236, 335–352 (2007).

39. Manni, L. et al. Ontology for the Asexual Development and Anatomy of the Colonial Chordate Botryllus schlosseri. PLoS ONE 9, e96434 (2014).

40. Rinkevich, B. The colonial urochordate Botryllus schlosseri: from stem cells and natural tissue transplantation to issues in evolutionary ecology. BioEssays 24, 730–740 (2002).

41. Tiozzo, S. et al. Embryonic versus blastogenetic development in the compound ascidian Botryllus schlosseri: Insights from Pitx expression patterns. Dev. Dyn. 232, 468–478 (2005).

42. Watanabe, H. Studies on the regulation in fused colonies in Botryllus primigenus (Ascidiae Compositae). Sci Rep Tokyo Bunrika Daigaku 7, 183–198 (1953).

43. Milkman, R. Genetic and developmental studies on Botryllus schlosseri. Biol. Bull. 132, 229–243 (1967).

44. Lauzon, R. J., Kidder, S. J. & Long, P. Suppression of programmed cell death regulates the cyclical degeneration of organs in a colonial urochordate. Dev. Biol. 301, 92–105 (2007).

45. Rinkevich, B. & Weissman, I. L. A long-term study on fused subclones in the ascidian Botryllus schlosseri: the resorption phenomenon (Protochordata: Tunicata). J. Zool. 213, 717–733 (1987).

46. Watterson, R. L. A sexual Reproduction in the Colonial Tunicate, Botryllus schlosseri (Pallas) Savigny, with Special Reference to the Developmental History of Intersiphonal Bands of Pigment Cells. Biol. Bull. 88, 71–103 (1945).

