## Supplementary material for "The *Orshina* rhythm in a colonial urochordate: recurrent aging/rejuvenation sequels": Suppl. Figures

**Supp. Fig 1.** Individual graphs for 35 NF colonies studied from birth to death. Observations were made every 15±5 days. Three parameters were followed: number of zooids, RS and CV. X axis shows the timescale from birth to death. Left y-axis shows the number of zooids. Right y-axis shows either RS or CV. Green curves = number of zooids. Red curves = RS. Black curves = CV. Black vertical lines are *Orshina* borders that mark the segments. Numbers above segments show lengths (days) of segments. Missing numbers represent cases where borders could not be set. These segments were not added to statistical analyses.

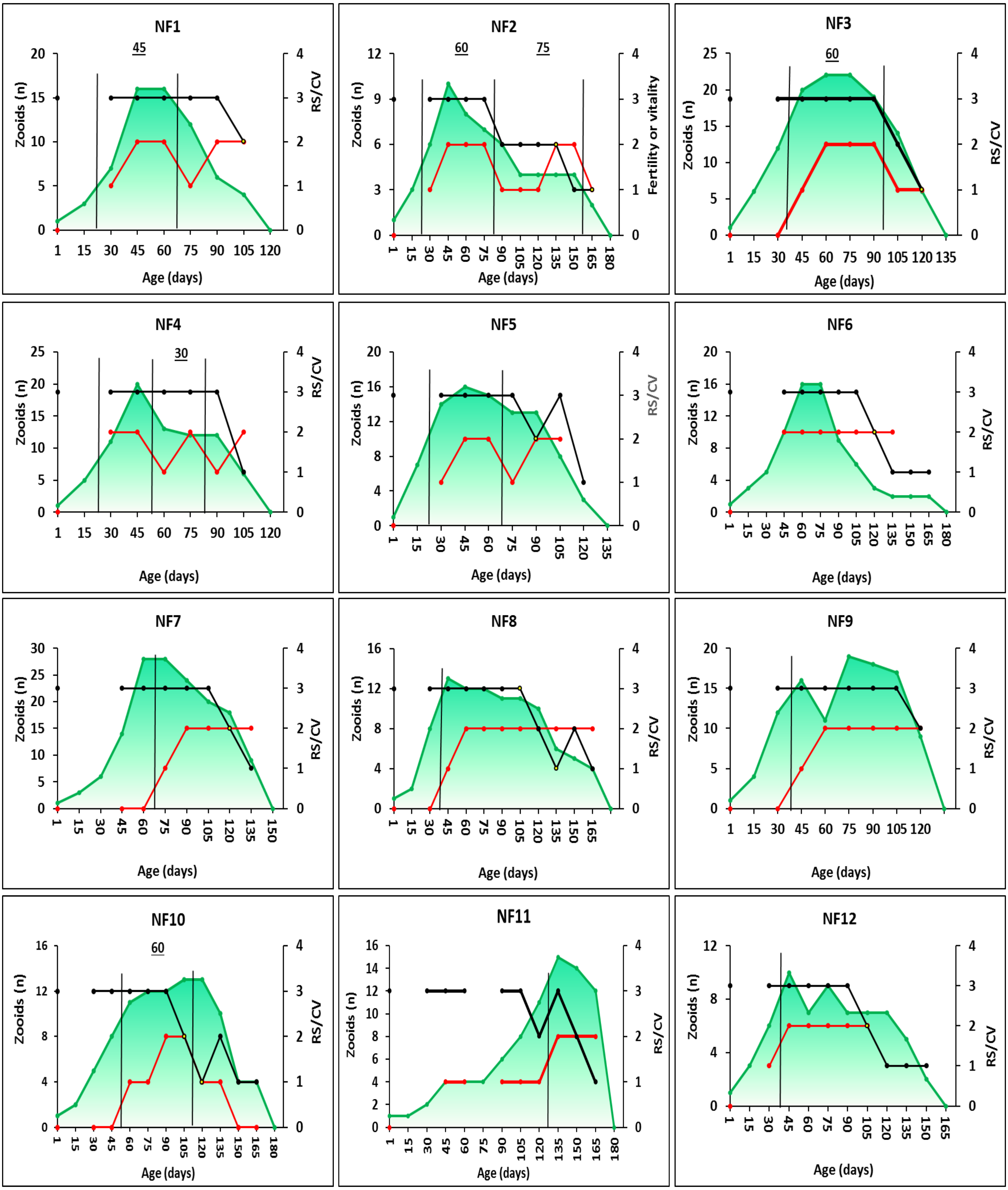

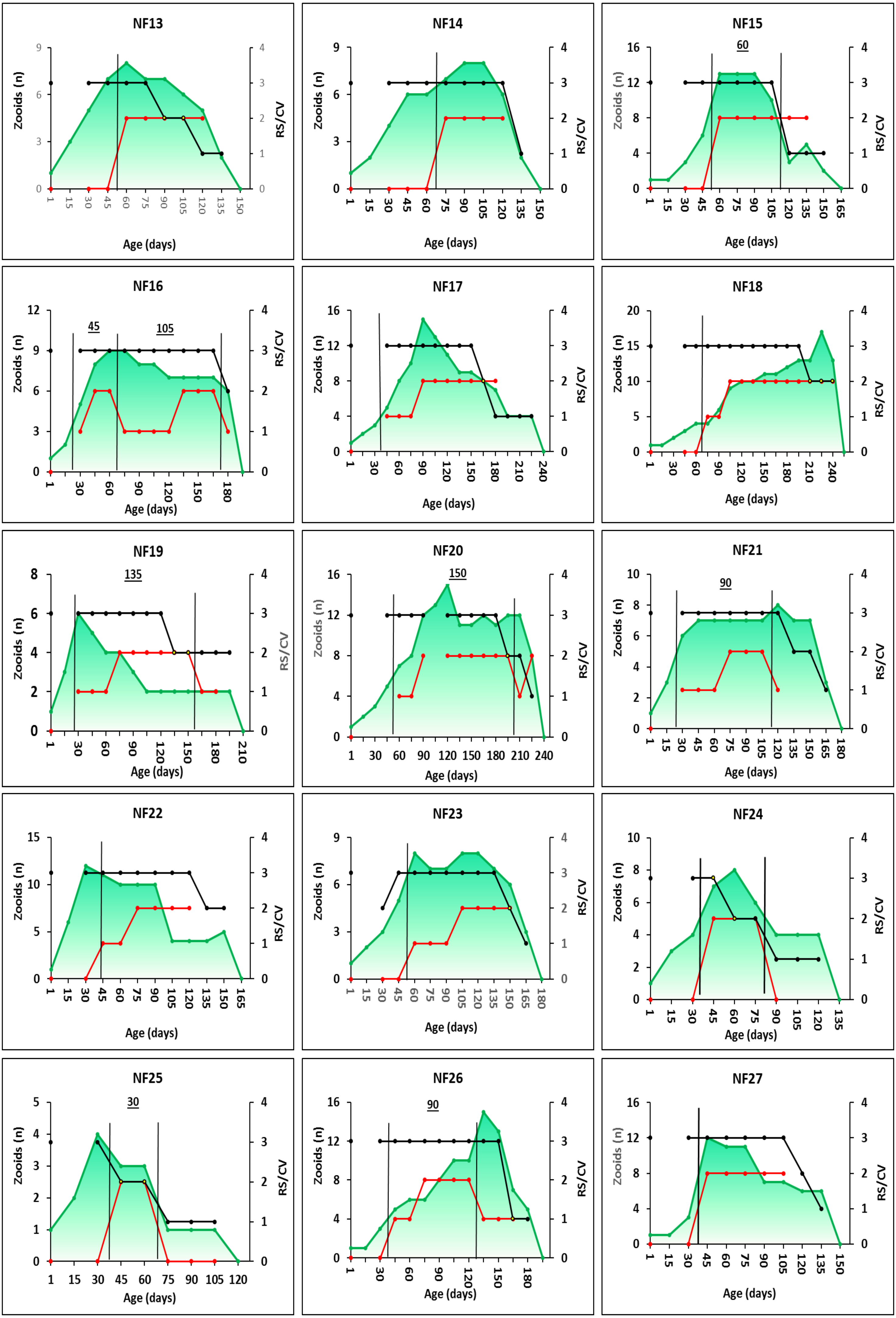
NF13
Zoids (n)
RS/CV
Age (days)
NF14
Zooids (n)
RS/CV
Age (days)
NF15
60
Zooids (n)
RS/CV
Age (days)
NF16
45
105
Zooids (n)
RS/CV
Age (days)
NF17
Zooids (n)
RS/CV
Age (days)
NF18
Zooids (n)
RS/CV
Age (days)
NF19
135
Zooids (n)
RS/CV
Age (days)
NF20
150
Zooids (n)
RS/CV
Age (days)
NF21
90
Zooids (n)
RS/CV
Age (days)
NF22
Zooids (n)
RS/CV
Age (days)
NF23
Zooids (n)
RS/CV
Age (days)
NF24
Zooids (n)
RS/CV
Age (days)
NF25
30
Zooids (n)
RS/CV
Age (days)
NF26
90
Zooids (n)
RS/CV
Age (days)
NF27
Zooids (n)
RS/CV
Age (days)

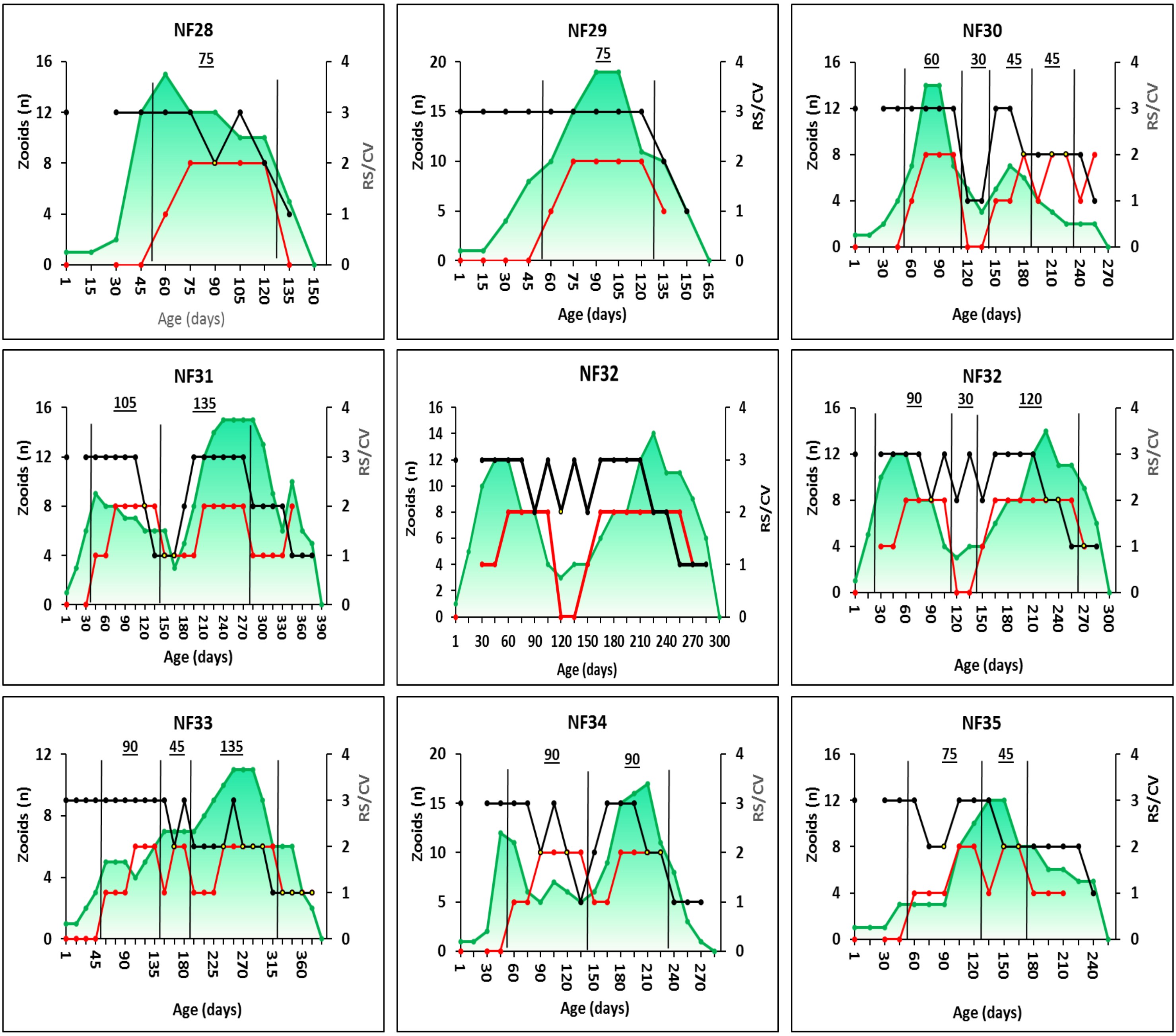
NF28
75
Zooids (n)
RS/CV
Age (days)
NF29
75
Zooids (n)
RS/CV
Age (days)
NF30
60
30
45
45
Zooids (n)
RS/CV
Age (days)
NF31
105
135
Zooids (n)
RS/CV
Age (days)
NF32
Zooids (n)
RS/CV
Age (days)
NF32
90
30
120
Zooids (n)
RS/CV
Age (days)
NF33
90
45
135
Zooids (n)
RS/CV
Age (days)
NF34
90
90
Zooids (n)
RS/CV
Age (days)
NF35
75
45
Zooids (n)
RS/CV
Age (days)

**Supp. Fig 2.** Individual graphs for 23 FA colonies studied from birth to death. Observations were made every 15±5 days. Three parameters were documented: number of zooids, RS and CV. X axis shows the timescale from birth to death. Left y-axis shows the number of zooids. Right y =axis shows either RS or CV. Green curves = number of zooids. Red curves = RS. Black curves = CV. Black vertical lines are *Orshina* borders that mark the segments. Numbers above segments show lengths (days) of segment. Fissions are marked with orange dashed lines. Deaths of ramets marked with red crosses. Missing numbers represent cases where borders could not be set. These segments were not added to statistical analyses.

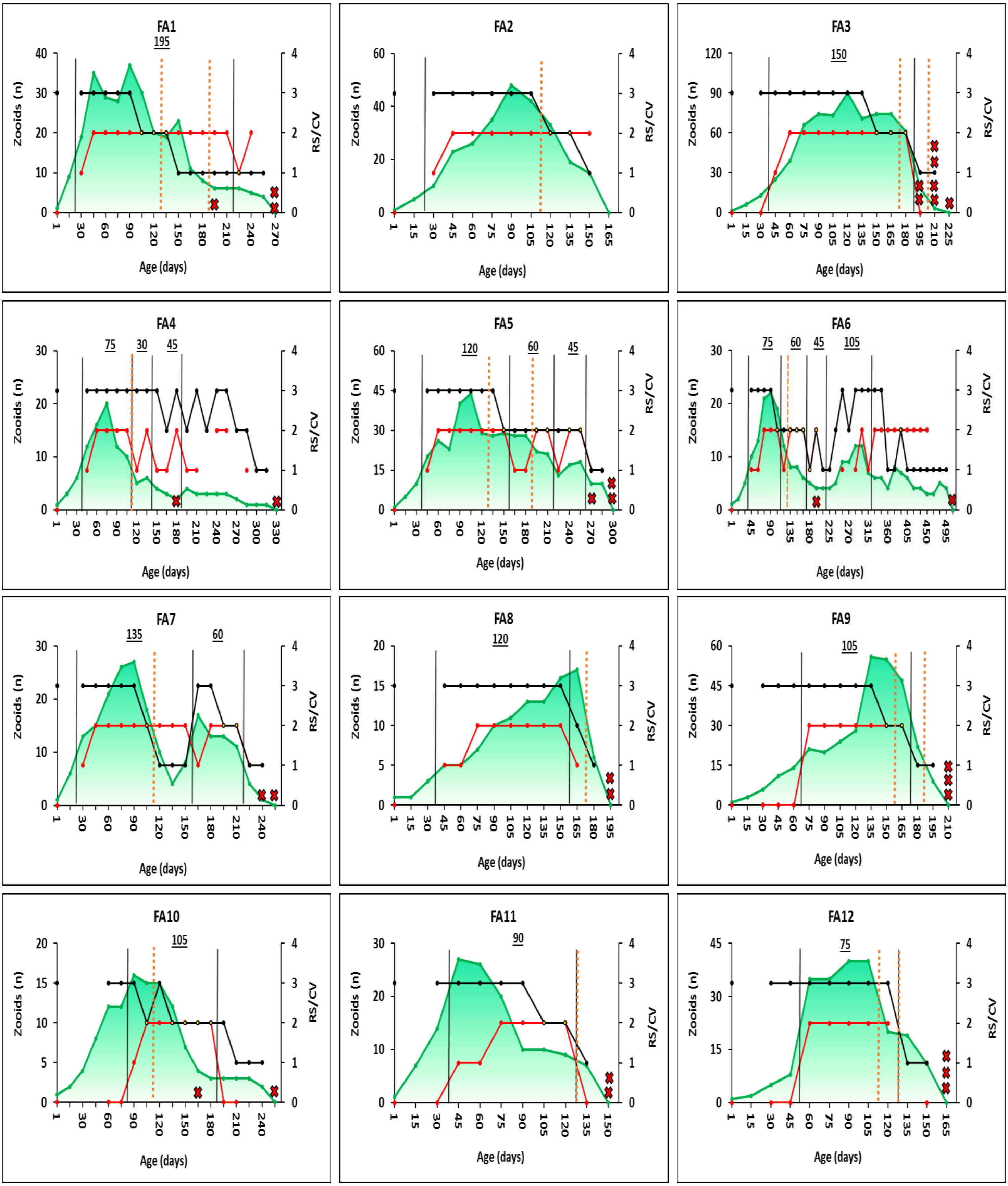

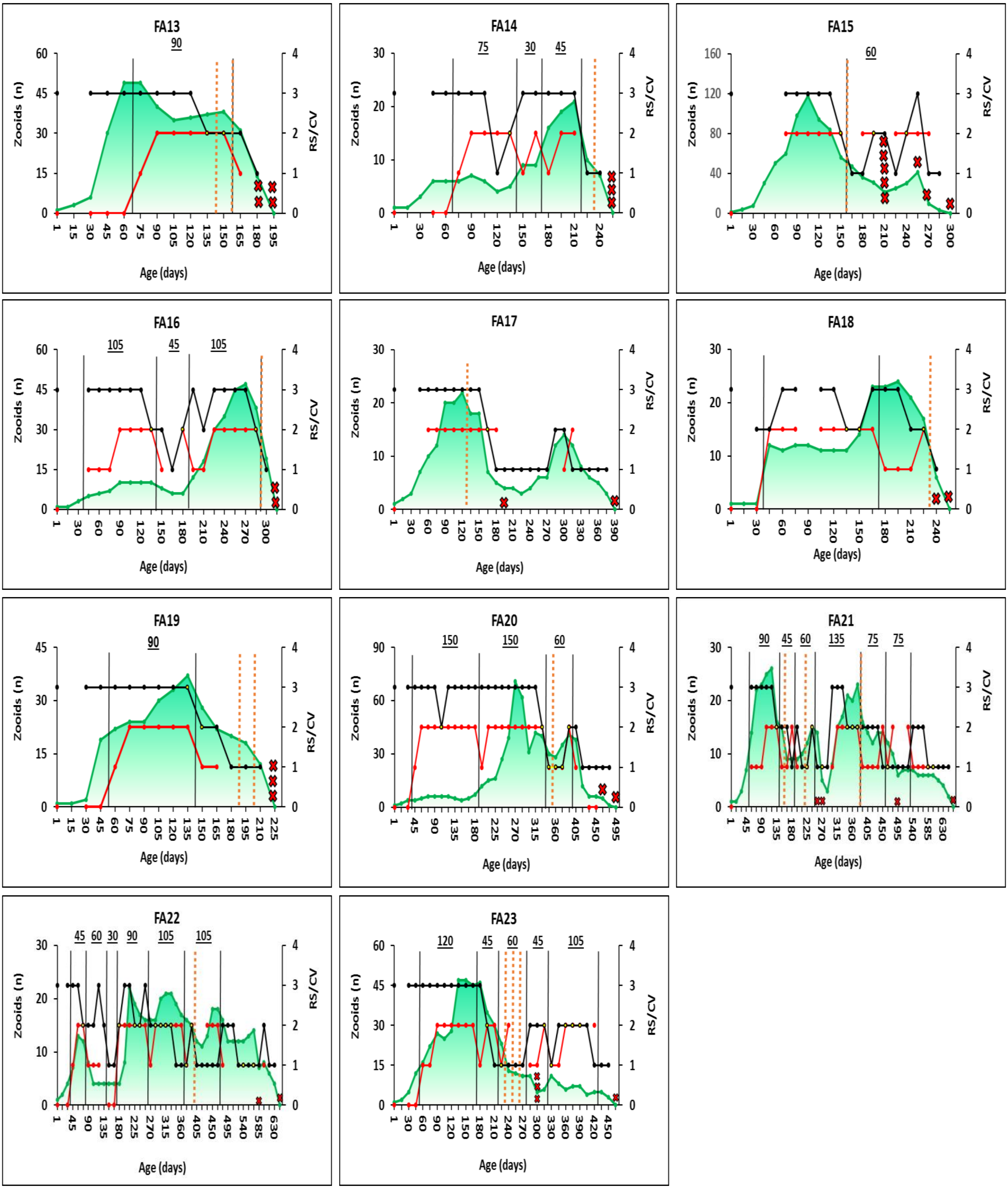
FA13
FA14
FA15
FA16
FA17
FA18
FA19
FA20
FA21
FA22
FA23
Zooids (n)
RS/CV
Age (days)

**Supp. Fig 3.** Individual graphs for 23 FB colonies studied from birth to death. Observations were made every 15±5 days. Three parameters were documented: number of zooids, RS and CV. X axis shows the timescale from birth to death. Left y-axis shows the number of zooids. Right y =axis shows either RS or CV. Green curves = number of zooids. Red curves = RS. Black curves = CV. Black vertical lines are *Orshina* borders that mark the segments. Numbers above segments show lengths (days) of segment. Fissions are marked with orange dashed lines. Deaths of ramets marked with red crosses. Missing numbers represent cases where borders could not be set. These segments were not added to statistical analyses.

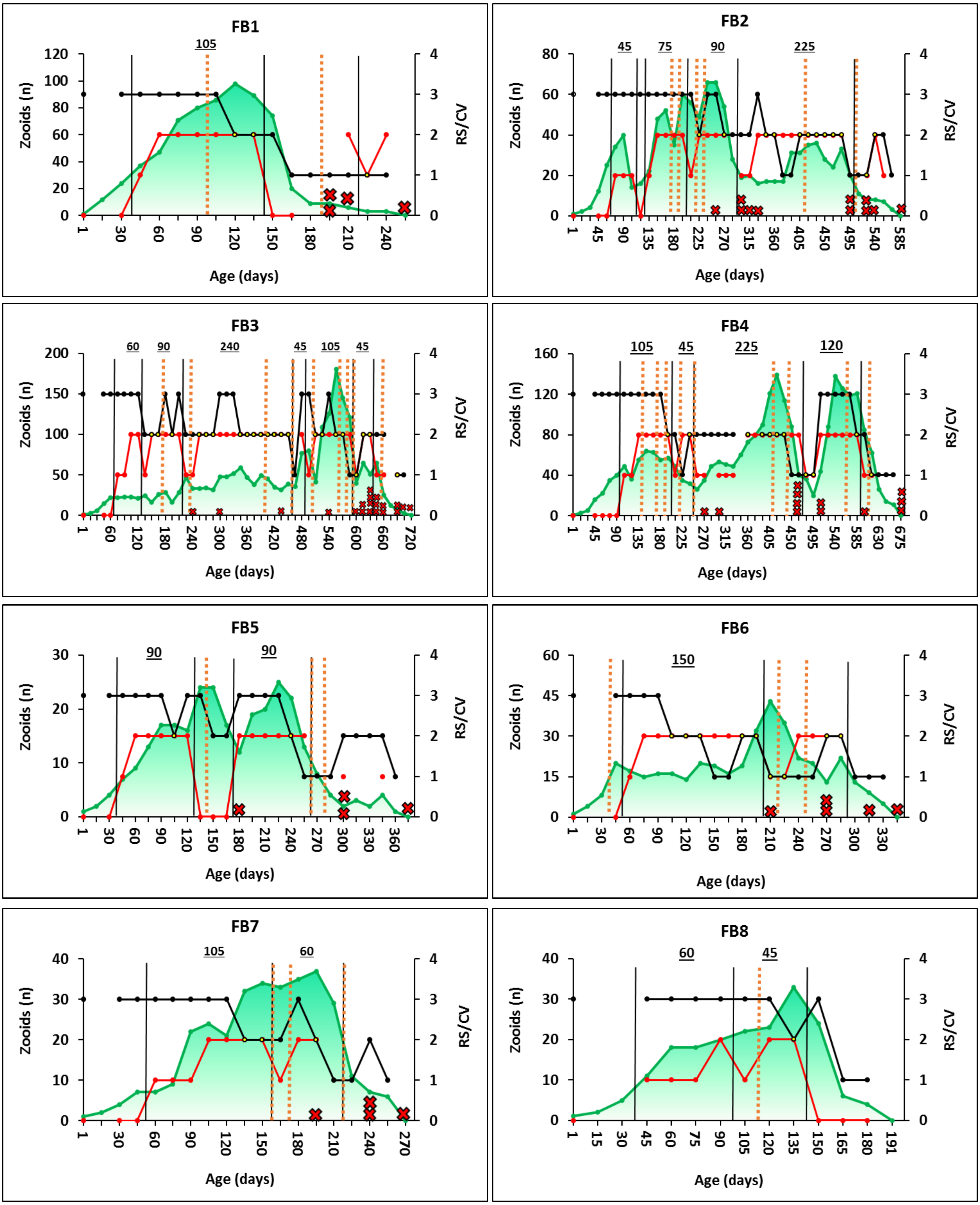

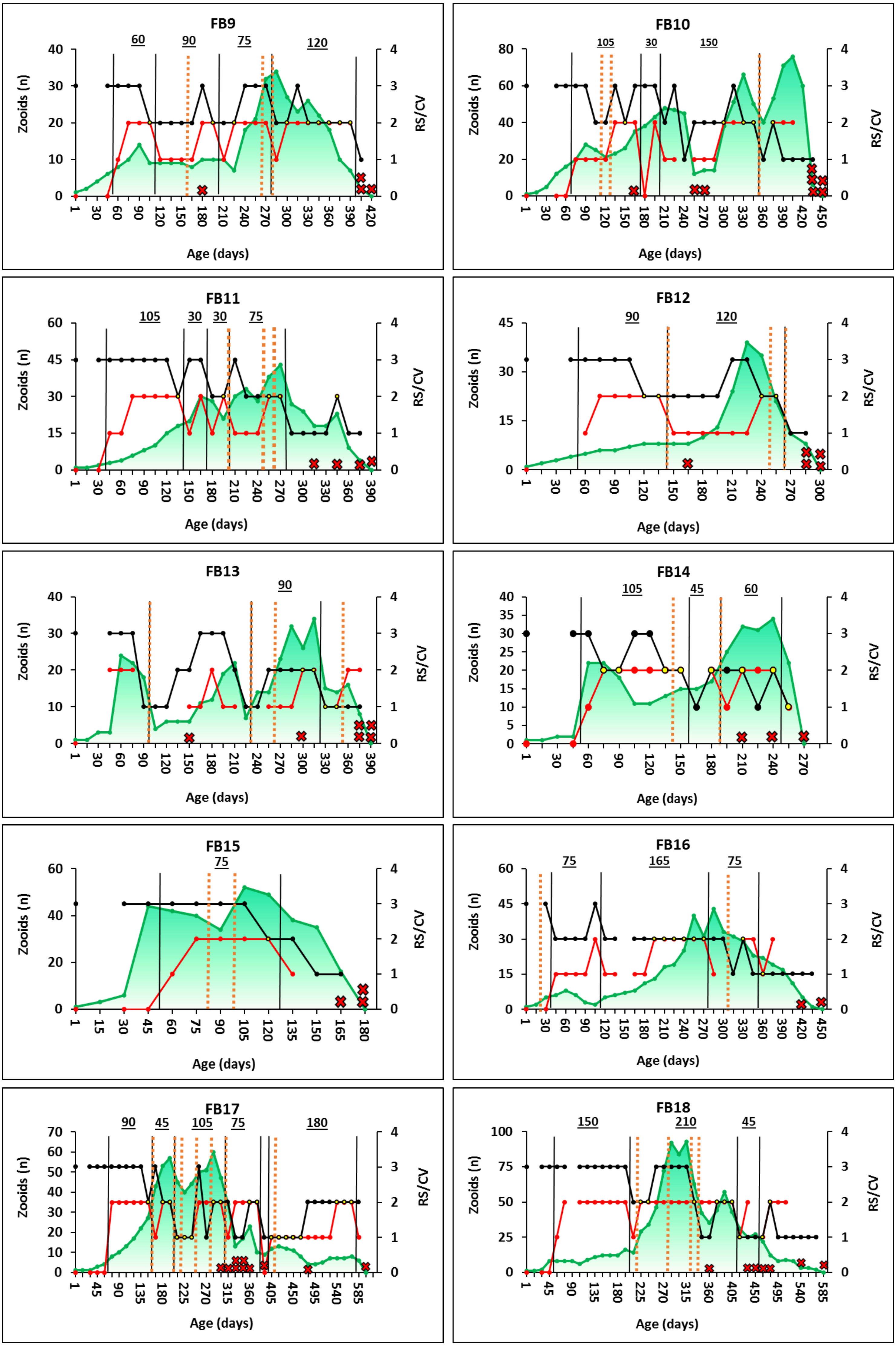
FB9
60
90
75
120
Zooids (n)
RS/CV
Age (days)
FB10
105
30
150
FB11
105
30
30
75
FB12
90
120
FB13
90
FB14
105
45
60
FB15
75
FB16
75
165
75
FB17
90
45
105
75
180
FB18
150
210
45

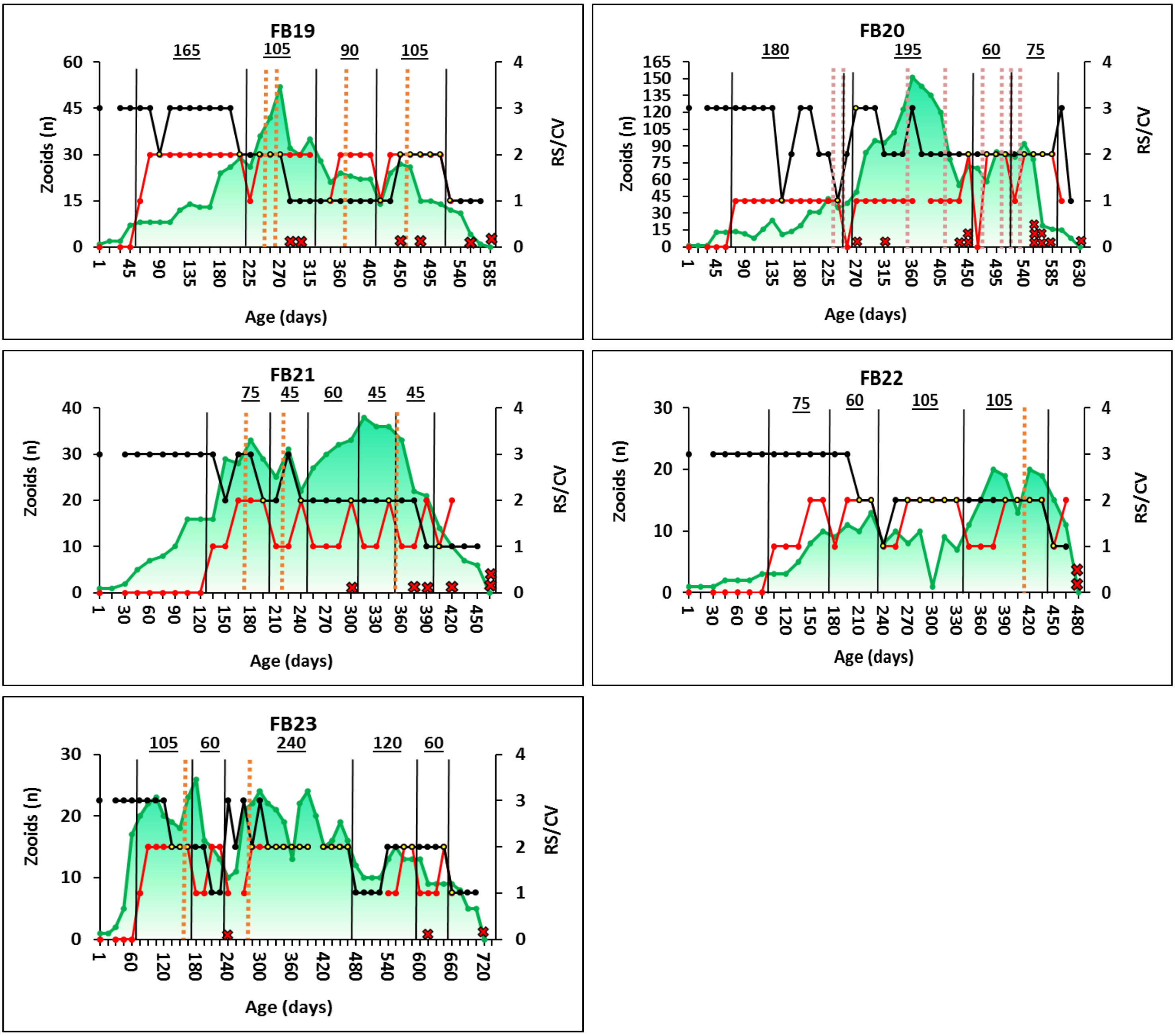

FB19
165
105
90
105
Zooids (n)
RS/CV
Age (days)
FB20
180
195
60
75
Zoods (n)
RS/CV
Age (days)
FB21
75
45
60
45
45
Zooids (n)
RS/CV
Age (days)
FB22
75
60
105
105
Zooids (n)
RS/CV
Age (days)
FB23
105
60
240
120
60
Zooids (n)
RS/CV
Age (days)
