## Supplementary material for "The *Orshina* rhythm in a colonial urochordate: recurrent aging/rejuvenation sequels": Suppl. Tables

**Suppl. Table 1.** Statistical outcomes for each of the three life history strategies: (a) Results of Spearman's correlations between pairs of variables (1st = Numbers of zooids; 2nd = RS; 3rd = CV). (b) Results of *p* values calculated using one-sample T-tests (against a "test value" of $r_s$ =0) for the correlations in Suppl. Table 1a, for each life-history type and for each pair of parameters.

**a**

| NF | Zooids vs. RS | Zooids vs. CV | RS vs. CV |
|---|---|---|---|
| NF1 | 0.35 | 0.36 | -0.34 |
| NF2 | 0.62 | 0.55 | 0.07 |
| NF3 | 0.81 | 0.41 | 0.13 |
| NF4 | 0.24 | 0.41 | -0.34 |
| NF5 | 0.48 | 0.35 | -0.34 |
| NF6 | 0.58 | 0.61 | -0.22 |
| NF7 | 0.04 | 0.41 | -0.55 |
| NF8 | 0.16 | 0.56 | -0.45 |
| NF9 | 0.45 | 0.41 | -0.29 |
| NF10 | 0.87 | -0.02 | 0.06 |
| NF11 | 0.90 | -0.56 | -0.55 |
| NF12 | 0.83 | 0.34 | -0.26 |
| NF13 | 0.47 | 0.33 | -0.59 |
| NF14 | 0.79 | 0.42 | ? |
| NF15 | 0.61 | 0.46 | -0.38 |
| NF16 | 0.35 | 0.31 | 0.19 |
| NF17 | 0.56 | 0.35 | -0.35 |
| NF18 | 0.83 | -0.66 | -0.35 |
| NF19 | -0.02 | 0.46 | -0.06 |
| NF20 | 0.57 | -0.18 | -0.06 |
| NF21 | 0.04 | 0.22 | ? |
| NF22 | -0.24 | 0.27 | ? |
| NF23 | 0.69 | 0.56 | 0.05 |
| NF24 | 0.89 | 0.07 | -0.11 |
| NF25 | 0.53 | 0.60 | 0.17 |
| NF26 | 0.51 | 0.16 | 0.11 |
| NF27 | 0.81 | 0.31 | ? |
| NF28 | 0.47 | 0.10 | -0.14 |
| NF29 | 0.94 | 0.13 | 0.00 |
| NF30 | 0.28 | 0.41 | 0.04 |
| NF31 | 0.43 | 0.42 | 0.14 |
| NF32 | 0.48 | -0.06 | 0.04 |
| NF33 | 0.64 | -0.27 | -0.14 |
| NF34 | 0.35 | 0.33 | -0.52 |
| NF35 | 0.78 | -0.14 | -0.24 |
| **Mean** | 0.52 | 0.24 | -0.17 |
| **STD** | 0.29 | 0.31 | 0.24 |

| FA | Zooids vs. RS | Zooids vs. CV | RS vs. CV |
|---|---|---|---|
| FA1 | 0.56 | 0.38 | -0.34 |
| FA2 | 0.79 | 0.09 | -0.37 |
| FA3 | 0.82 | 0.13 | 0.03 |
| FA4 | 0.43 | 0.59 | 0.41 |
| FA5 | 0.55 | 0.20 | 0.04 |
| FA6 | -0.11 | 0.62 | -0.40 |
| FA7 | 0.24 | 0.64 | -0.43 |
| FA8 | 0.50 | -0.18 | 0.33 |
| FA9 | 0.84 | -0.18 | -0.36 |
| FA10 | 0.34 | 0.51 | -0.28 |
| FA11 | 0.22 | 0.53 | -0.08 |
| FA12 | 0.87 | 0.18 | 0.40 |
| FA13 | 0.39 | 0.11 | -0.38 |
| FA14 | 0.29 | 0.20 | -0.38 |
| FA15 | 0.46 | 0.56 | -0.26 |
| FA16 | 0.59 | 0.04 | -0.05 |
| FA17 | 0.36 | 0.64 | 0.05 |
| FA18 | 0.06 | 0.20 | -0.02 |
| FA19 | 0.86 | 0.29 | 0.16 |
| FA20 | 0.27 | -0.07 | 0.50 |
| FA21 | 0.52 | 0.56 | -0.05 |
| FA22 | 0.68 | -0.06 | 0.01 |
| FA23 | 0.43 | 0.29 | -0.24 |
| **Mean** | 0.48 | 0.27 | -0.07 |
| **STD** | 0.26 | 0.27 | 0.29 |

| FB | Zooids vs. RS | Zooids vs. CV | RS vs. CV |
|---|---|---|---|
| FB1 | 0.45 | 0.45 | 0.02 |
| FB2 | 0.53 | 0.35 | -0.30 |
| FB3 | 0.40 | -0.09 | -0.12 |
| FB4 | 0.68 | 0.18 | -0.09 |
| FB5 | 0.24 | 0.31 | 0.14 |
| FB6 | -0.14 | -0.16 | -0.26 |
| FB7 | 0.80 | -0.24 | -0.43 |
| FB8 | 0.88 | 0.23 | 0.13 |
| FB9 | 0.43 | 0.13 | 0.14 |
| FB10 | 0.68 | -0.33 | 0.37 |
| FB11 | 0.32 | -0.30 | -0.23 |
| FB12 | 0.13 | -0.42 | -0.18 |
| FB13 | 0.25 | 0.10 | 0.04 |
| FB14 | 0.28 | -0.37 | -0.34 |
| FB15 | 0.49 | 0.09 | -0.22 |
| FB16 | 0.58 | -0.25 | -0.34 |
| FB17 | 0.61 | -0.27 | -0.04 |
| FB18 | 0.42 | 0.05 | 0.05 |
| FB19 | 0.49 | -0.31 | -0.12 |
| FB20 | 0.40 | -0.32 | -0.41 |
| FB21 | 0.59 | -0.18 | -0.59 |
| FB22 | 0.52 | -0.60 | -0.47 |
| FB23 | 0.48 | 0.34 | -0.33 |
| **Mean** | 0.46 | -0.07 | -0.16 |
| **STD** | 0.22 | 0.29 | 0.24 |

**b**

| Life history type | Zooids vs. RS | Zooids vs. CV | RS vs. CV |
|---|---|---|---|
| **NF** | <0.001 | <0.001 | <0.001 |
| **FA** | <0.001 | <0.001 | NS |
| **FB** | <0.001 | NS | 0.005 |

**Suppl. Table 2.** Reproductive status along RS segments. In each segment, the number of time RS male and RS female was counted. Some of the colonies have more than one RS segment, therefore they are exhibited in more than one line. All life history types were accounted for this analysis. The counts were converted into percent where 100% is the total RS statuses counted in each RS segment. In certain places data is missing due to inability to determine RS borders in those animals.

| NF | Number of time Testes counted in RS segment | Number of time ova counted in RS segment | Sum of all ova and testes counted in a RS segment | %T | %O |
|---|---|---|---|---|---|
| NF1 | 1 | 2 | 3 | 33 | 67 |
| NF2 | 1 | 3 | 4 | 25 | 75 |
| NF2 | 3 | 2 | 5 | 60 | 40 |
| NF3 | 1 | 3 | 4 | 25 | 75 |
| NF4 | 1 | 1 | 2 | 50 | 50 |
| NF5 | 1 | 2 | 3 | 33 | 67 |
| NF6 | ? | ? | | | |
| NF7 | ? | ? | | | |
| NF8 | ? | ? | | | |
| NF9 | ? | ? | | | |
| NF10 | 2 | 2 | 4 | 50 | 50 |
| NF11 | ? | ? | | | |
| NF12 | ? | ? | | | |
| NF13 | ? | ? | | | |
| NF14 | ? | ? | | | |
| NF15 | 0 | 4 | 4 | 0 | 100 |
| NF16 | 1 | 2 | 3 | 33 | 67 |
| NF16 | 4 | 3 | 7 | 57 | 43 |
| NF17 | ? | ? | | | |
| NF18 | ? | ? | | | |
| NF19 | 2 | 7 | 9 | 22 | 78 |
| NF20 | ? | ? | | | |
| NF21 | 3 | 3 | 6 | 50 | 50 |
| NF22 | ? | ? | | | |
| NF23 | ? | ? | | | |
| NF24 | 0 | 3 | 3 | 0 | 100 |
| NF25 | ? | ? | | | |
| NF26 | 2 | 4 | 6 | 33 | 67 |
| NF27 | ? | ? | | | |
| NF28 | 1 | 4 | 5 | 20 | 80 |
| NF29 | 1 | 4 | 5 | 20 | 80 |
| NF30 | 1 | 3 | 4 | 25 | 75 |
| NF30 | 2 | 1 | 3 | 67 | 33 |
| NF30 | 1 | 2 | 3 | 33 | 67 |
| NF31 | 2 | 5 | 7 | 29 | 71 |
| NF31 | 4 | 5 | 9 | 44 | 56 |
| NF32 | 2 | 4 | 6 | 33 | 67 |
| NF32 | 1 | 7 | 8 | 13 | 88 |
| NF33 | 3 | 3 | 6 | 50 | 50 |
| NF33 | 1 | 2 | 3 | 33 | 67 |
| NF33 | 3 | 6 | 9 | 33 | 67 |
| NF34 | 2 | 4 | 6 | 33 | 67 |
| NF34 | 2 | 4 | 6 | 33 | 67 |
| NF35 | 3 | 2 | 5 | 60 | 40 |
| NF35 | 1 | 2 | 3 | 33 | 67 |
| Mean | 1.7 | 3.3 | | 34 | 66 |
| STD | 1.0 | 1.6 | | 16 | 16 |
| n | 30 | 30 | | | |

| FA | Number of time Testes counted in RS segment | Number of time ova counted in RS segment | Sum of all ova and testes counted in a RS segment | %T | %O |
|---|---|---|---|---|---|
| FA1 | ? | ? | | | |
| FA2 | ? | ? | | | |
| FA3 | 1 | 9 | 10 | 10 | 90 |
| FA4 | 1 | 4 | 5 | 20 | 80 |
| FA4 | 1 | 1 | 2 | 50 | 50 |
| FA4 | 2 | 1 | 3 | 67 | 33 |
| FA5 | 1 | 5 | 6 | 17 | 83 |
| FA5 | 2 | 2 | 4 | 50 | 50 |
| FA5 | 1 | 2 | 3 | 33 | 67 |
| FA6 | 2 | 3 | 5 | 40 | 60 |
| FA6 | 1 | 3 | 4 | 25 | 75 |
| FA7 | 1 | 8 | 9 | 11 | 89 |
| FA8 | 2 | 6 | 8 | 25 | 75 |
| FA9 | 0 | 9 | 9 | 0 | 100 |
| FA10 | 1 | 6 | 7 | 14 | 86 |
| FA11 | 2 | 4 | 6 | 33 | 67 |
| FA12 | ? | ? | | | |
| FA13 | 1 | 5 | 6 | 17 | 83 |
| FA14 | 1 | 4 | 5 | 20 | 80 |
| FA14 | 1 | 1 | 2 | 50 | 50 |
| FA15 | ? | ? | | | |
| FA16 | 3 | 4 | 7 | 43 | 57 |
| FA16 | 2 | 5 | 7 | 29 | 71 |
| FA17 | ? | ? | | | |
| FA18 | ? | ? | | | |
| FA19 | 1 | 5 | 6 | 17 | 83 |
| FA20 | 1 | 9 | 10 | 10 | 90 |
| FA20 | 1 | 9 | 10 | 10 | 90 |
| FA20 | 3 | 1 | 4 | 75 | 25 |
| FA21 | 3 | 3 | 6 | 50 | 50 |
| FA21 | 2 | 1 | 3 | 67 | 33 |
| FA21 | 4 | 1 | 5 | 80 | 20 |
| FA22 | 1 | 2 | 3 | 33 | 67 |
| FA22 | 1 | 6 | 7 | 14 | 86 |
| FA23 | 2 | 6 | 8 | 25 | 75 |
| FA23 | 1 | 2 | 3 | 33 | 67 |
| FA23 | 2 | 1 | 3 | 67 | 33 |
| Mean | 1.5 | 4.1 | | 33 | 67 |
| STD | 0.9 | 2.7 | | 22 | 22 |
| n | 31 | 31 | | | |

| FB | Number of time Testes counted in RS segment | Number of time ova counted in RS segment | Sum of all ova and testes counted in a RS segment | %T | %O |
|---|---|---|---|---|---|
| FB1 | 1 | 6 | 7 | 14 | 86 |
| FB2 | 1 | 4 | 5 | 20 | 80 |
| FB3 | 2 | 2 | 4 | 50 | 50 |
| FB3 | 1 | 5 | 6 | 17 | 83 |
| FB3 | 2 | 14 | 16 | 13 | 88 |
| FB3 | 1 | 6 | 7 | 14 | 86 |
| FB3 | 1 | 2 | 3 | 33 | 67 |
| FB4 | 2 | 5 | 7 | 29 | 71 |
| FB4 | 1 | 2 | 3 | 33 | 67 |
| FB4 | 2 | 6 | 8 | 25 | 75 |
| FB5 | 2 | 4 | 6 | 33 | 67 |
| FB6 | 1 | 9 | 10 | 10 | 90 |
| FB7 | 3 | 4 | 7 | 43 | 57 |
| FB8 | 1 | 2 | 3 | 33 | 67 |
| FB9 | 1 | 3 | 4 | 25 | 75 |
| FB9 | 4 | 2 | 6 | 67 | 33 |
| FB9 | 1 | 4 | 5 | 20 | 80 |
| FB10 | 4 | 3 | 7 | 57 | 43 |
| FB11 | 2 | 5 | 7 | 29 | 71 |
| FB11 | 1 | 1 | 2 | 50 | 50 |
| FB11 | 1 | 1 | 2 | 50 | 50 |
| FB11 | 3 | 2 | 5 | 60 | 40 |
| FB12 | 1 | 5 | 6 | 17 | 83 |
| FB13 | ? | | | | |
| FB14 | 1 | 6 | 7 | 14 | 86 |
| FB14 | 1 | 3 | 4 | 25 | 75 |
| FB15 | 1 | 4 | 5 | 20 | 80 |
| FB16 | 4 | 1 | 5 | 80 | 20 |
| FB17 | 1 | 2 | 3 | 33 | 67 |
| FB17 | 3 | 4 | 7 | 43 | 57 |
| FB17 | 1 | 4 | 5 | 20 | 80 |
| FB17 | 9 | 3 | 12 | 75 | 25 |
| FB18 | 1 | 13 | 14 | 7 | 93 |
| FB19 | 1 | 1 | 2 | 50 | 50 |
| FB19 | 1 | 6 | 7 | 14 | 86 |
| FB20 | ? | | | | |
| FB21 | 2 | 3 | 5 | 40 | 60 |
| FB21 | 2 | 1 | 3 | 67 | 33 |
| FB21 | 3 | 1 | 4 | 75 | 25 |
| FB21 | 2 | 1 | 3 | 67 | 33 |
| FB21 | 2 | 1 | 3 | 67 | 33 |
| FB22 | 3 | 2 | 5 | 60 | 40 |
| FB22 | 1 | 3 | 4 | 25 | 75 |
| FB22 | 2 | 5 | 7 | 29 | 71 |
| FB22 | 3 | 4 | 7 | 43 | 57 |
| FB23 | 1 | 6 | 7 | 14 | 86 |
| FB23 | 2 | 2 | 4 | 50 | 50 |
| Mean | 1.9 | 3.8 | | 37 | 63 |
| STD | 1.4 | 2.8 | | 20 | 20 |
| n | 45 | 45 | | | |

**Suppl. Table 3.** Simple cases of colonies, where fission occurred once. The total life span of the ramets (days) is shown. Chi square test is described in Suppl. Table 4.

| | Name | 1st Ramet death | 2nd Ramet death |
|---|---|---|---|
| **1** | FA4 | 180 | 330 |
| **2** | FA6 | 195 | 510 |
| **3** | FA7 | 240 | 255 |
| **4** | FA8 | 195 | 195 |
| **5** | FA10 | 165 | 255 |
| **6** | FA11 | 150 | 150 |
| **7** | FA16 | 315 | 315 |
| **8** | FA17 | 195 | 390 |
| **9** | FA18 | 240 | 255 |
| **10** | FA19 | 225 | 225 |
| **11** | FA20 | 465 | 495 |
| **12** | FA22 | 645 | 585 |
| **13** | FB16 | 420 | 450 |
| **14** | FB22 | 480 | 480 |

**Suppl. Table 4.** Chi-square test designed to evaluate whether ramets die at the same time (e.g., experience nonrandom deaths, performed on cases with a single fission events). (a) Schematic illustration demonstrating the reasoning for selecting expected probabilities to be used in the following chi-test. The relevant timeline for ramets' death is from the first fission event to death of the genet. The timeline is marked by a double arrowheads line and equals 208 days; the mean age of deaths in the FA and FB life history strategies minus the mean age for the complete first fission event in both strategies). The timeline (=208d) is composed of 14 observations (15 days, each). Accordingly, the expected probability of two ramets to be found dead in the same observation is 1/14 (=0.07). This probability was used in the first chi-square test. (b) Summary table for the probabilities and test results.

a

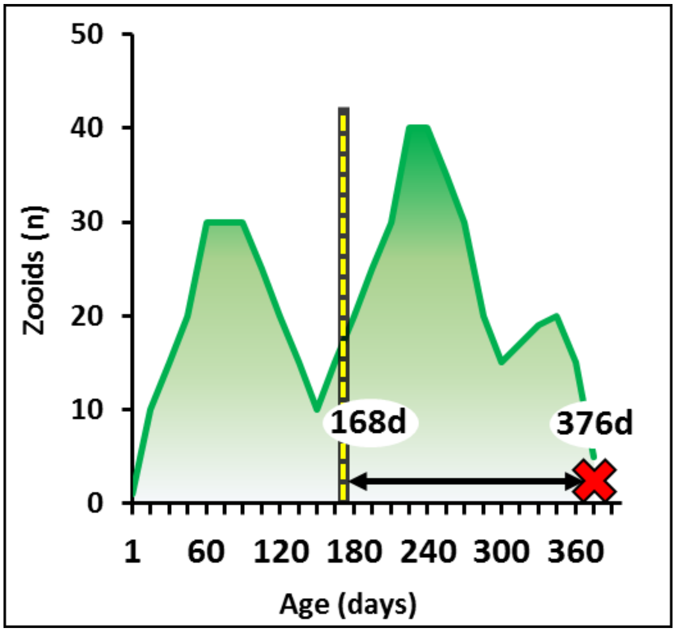


b

| | Hypothesized proportion | Observed | Expected | Chi Square |
|---|---|---|---|---|
| **Death of ramets is at same observation** | 0.07 | 5 | 0.98 | 16.49 |
| **Death of ramets is not at same observation** | 0.93 | 9 | 13.02 | 1.24 |
| | | 14 | | 17.73, df=1, *p*<0.0001 |

**Suppl. Table 5.** Chi-square test designed to evaluate whether ramets found dead significantly adjacent to segments borders or not. (a) A schematic illustration for the reasoning of selecting expected probabilities in the following test, based on the mean time for an Orshina segment (marked with a double arrowheads line. The length of each such segment is highlighted by either blue or red zones. The red zones (the two sides from each Orshina border), signify the time considered as ‘adjacent’ to the Orshina segments’ borders, each equal to 15 days (observation intervals). The blue zones signify the time considered as ‘away’ from the borders, equals 78 days. Accordingly, the expected probability of a ramet to die in border region (red zones) is 15/93. This probability was used in the following chi-square test. (b) Summary table for colonies participated in the test (n=29). The table signifies ramets dying adjacent to the border and ramets that die away from the Orshina borders. The table refer to Suppl. Figs. 2-3. (c) Summary table for the probabilities and test results.

**a**

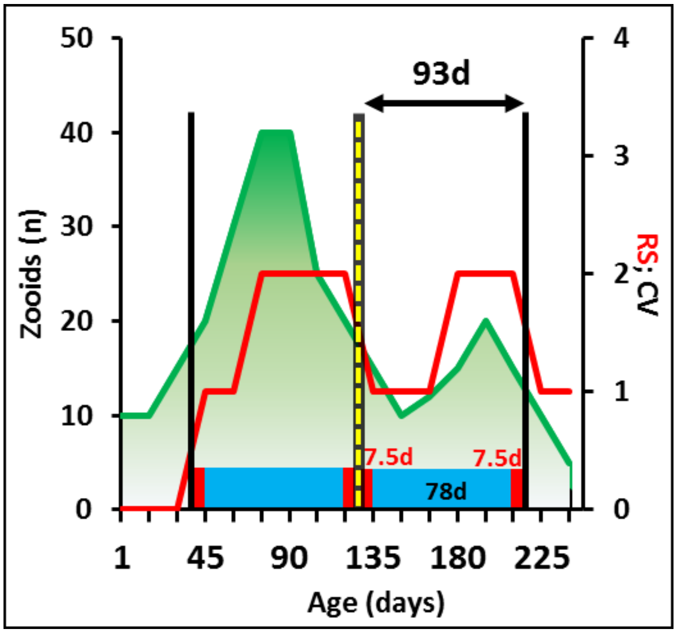


**b**

| | Adjacent | Non-adjacent |
|---|---|---|
| **FA1** | 0 | 1 |
| **FA3** | 2 | 5 |
| **FA4** | 1 | 0 |
| **FA5** | 1 | 2 |
| **FA6** | 0 | 1 |
| **FA7** | 0 | 2 |
| **FA10** | 0 | 1 |
| **FA15** | 5 | 0 |
| **FA16** | 0 | 2 |
| **FA21** | 1 | 2 |
| **FA23** | 0 | 3 |
| **FB1** | 1 | 2 |
| **FB2** | 4 | 7 |
| **FB3** | 7 | 8 |
| **FB4** | 5 | 4 |
| **FB5** | 1 | 2 |
| **FB6** | 1 | 2 |
| **FB7** | 0 | 3 |
| **FB9** | 2 | 2 |
| **FB10** | 1 | 2 |
| **FB12** | 0 | 3 |
| **FB13** | 0 | 2 |
| **FB14** | 1 | 2 |
| **FB17** | 3 | 6 |
| **FB18** | 2 | 3 |
| **FB19** | 0 | 4 |
| **FB20** | 4 | 7 |
| **FB21** | 2 | 2 |
| **FB23** | 1 | 1 |
| | 45 | 81 |

**c**

| | **Hypothesized proportion** | **Observed** | **Expected** | **Chi Square** |
|---|---|---|---|---|
| **Deaths are adjacent** | 0.16 | 45 | 20.16 | 30.6 |
| **Deaths are not adjacent** | 0.84 | 81 | 105.84 | 5.8 |
| | | 126 | 126 | 36.4, df=1, $p$<0.0001 |

**Suppl. Table 6.** Chi square test designed to evaluate whether fission events occur significantly adjacent to segments borders or not. (a) A schematic illustration for the reasoning of selecting expected probabilities to be used in the following test, based on the mean time for an Orshina segment (marked with a double arrowheads line.The length of each such segment is highlighted by either blue or red zones. The red zones (the two sides from each Orshina border), signify the time considered as ‘adjacent’ to the Orshina segments’ borders, each equal to 30 days (observation intervals). The blue zones signify the time considered as ‘away’ from the borders, equals 63 days. Accordingly, the expected probability of a ramet to die in border region (red zones) is 30/93. This probability was used in the following chi-square test. (b) Summary table for colonies participated in the test (n=41). The table signifies fission events adjacent to the border and fission events away from the borders. The table refer to Suppl. Figs. 2-3. (c) Summary table for the probabilities and test results.

**a**

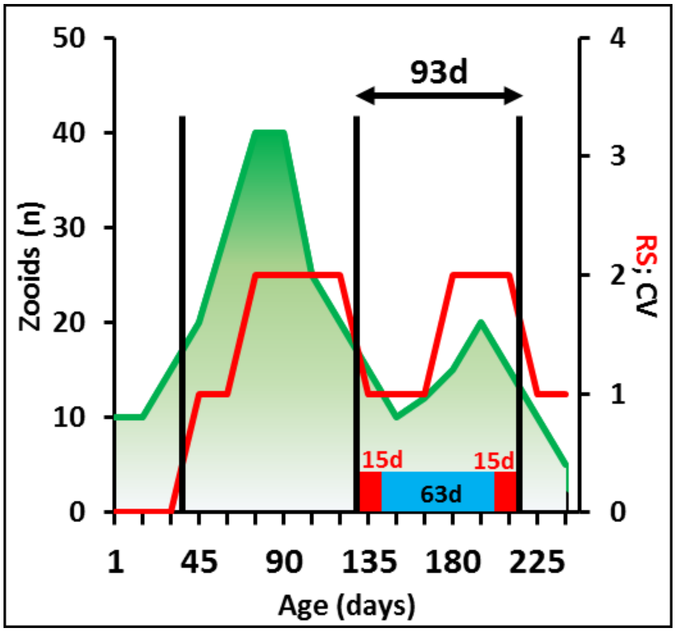


**b**

| Name | Adjacent | Non-adjacent |
|---|---|---|
| FA1 | 0 | 5 |
| FA3 | 6 | 0 |
| FA4 | 1 | 0 |
| FA5 | 0 | 2 |
| FA6 | 1 | 0 |
| FA7 | 0 | 1 |
| FA8 | 1 | 0 |
| FA9 | 2 | 0 |
| FA10 | 0 | 1 |
| FA11 | 1 | 0 |
| FA12 | 2 | 0 |
| FA13 | 3 | 0 |
| FA14 | 2 | 0 |
| FA15 | 7 | 0 |
| FA16 | 1 | 0 |
| FA20 | 1 | 0 |
| FA21 | 2 | 1 |
| FA22 | 0 | 1 |
| FA23 | 2 | 1 |
| FB2 | 5 | 3 |
| FB3 | 12 | 4 |
| FB4 | 4 | 6 |
| FB5 | 3 | 0 |
| FB6 | 2 | 2 |
| FB7 | 3 | 0 |
| FB8 | 1 | 0 |
| FB9 | 2 | 1 |
| FB10 | 4 | 3 |
| FB11 | 2 | 1 |
| FB12 | 4 | 0 |
| FB13 | 2 | 3 |
| FB14 | 2 | 0 |
| FB15 | 0 | 2 |
| FB16 | 1 | 1 |
| FB17 | 9 | 3 |
| FB18 | 1 | 6 |
| FB19 | 0 | 6 |
| FB20 | 10 | 4 |
| FB21 | 5 | 1 |
| FB22 | 0 | 1 |
| FB23 | 1 | 1 |
| Total | 105 | 60 |

**c**

| | Hypothesized proportion | Observed | Expected | Chi Square |
|---|---|---|---|---|
| Fission adjacent | 0.32 | 105 | 52.8 | 51.61 |
| Fission is not adjacent | 0.68 | 60 | 112.2 | 24.29 |
| | | 165 | 165 | 75.89, df=1, *p*<0.0001 |
